# Detection of naturally acquired, strain-transcending antibodies against rosetting *Plasmodium falciparum* strains in humans

**DOI:** 10.1101/2024.01.11.575252

**Authors:** Florence E. McLean, Yvonne Azasi, Cameron Sutherland, Emmanuel Toboh, Daniel Ansong, Tsiri Agbenyega, Gordon Awandare, J. Alexandra Rowe

**Affiliations:** Institute of Immunology and Infection Research, School of Biological Sciences, University of Edinburgh, UK; Manhyia District Hospital, Kumasi, Ghana; Kwame Nkrumah University of Science and Technology, School of Medical Sciences, Kumasi, Ghana, Departments of Child Health and Medicine, Komfo Anokye Teaching Hospital, Kumasi, Ghana; Malaria Research Centre, Agogo, Ghana; West African Centre for Cell Biology of Infectious Pathogens, University of Ghana, Legon, Ghana

**Keywords:** severe malaria, PfEMP1, rosette formation, virulence, immunology, antibodies, epitopes

## Abstract

Strain-transcending antibodies against subsets of *P. falciparum* blood stage surface antigens could protect children from severe malaria. However, the evidence supporting the existence of such antibodies is incomplete and inconsistent. One subset of surface antigens associated with severe malaria, rosette-mediating *Plasmodium falciparum* Erythrocyte Membrane Protein one (PfEMP1) variants, cause infected erythrocytes to bind to uninfected erythrocytes to form clusters of cells (rosettes) that contribute to microvascular obstruction and pathology. Here, using flow cytometry of live infected erythrocytes, we tested plasma from 80 individuals living in malaria-endemic regions for IgG recognition of the surface of four *P. falciparum* rosetting strains. Broadly-reactive plasma samples were then used in antibody elution experiments in which intact IgG was eluted from the surface of infected erythrocytes and transferred to heterologous rosetting strains to look for strain-transcending antibodies. We found that seroprevalence (percentage of samples that recognised each strain) against allopatric rosetting strains was high in adults (58% - 93%), but lower in children (5%-30%). Strain-transcending antibodies were present in nine out of eleven eluted antibody experiments, with six of these recognising multiple heterologous rosetting parasite strains. One eluate had rosette disrupting activity against heterologous strains, suggesting PfEMP1 as the likely target of the strain-transcending antibodies. Naturally acquired strain-transcending antibodies to rosetting *P. falciparum* strains in humans have not been directly demonstrated previously. Their existence suggests that such antibodies could play a role in clinical protection and raises the possibility that conserved epitopes recognised by strain-transcending antibodies could be targeted therapeutically.

## Introduction

During the blood stage of *Plasmodium falciparum* malaria, the parasite modifies the surface of its host erythrocyte by displaying variant surface antigens (VSAs) on the infected cell surface. These VSAs are derived from polymorphic protein families, the most well studied of which is *Plasmodium falciparum* erythrocyte membrane protein 1 (PfEMP1), which is a family of adhesion proteins that mediates sequestration of infected cells in the microvasculature [1]. Other VSA families include the repetitive interspersed family (RIFIN), subtelomeric variant open reading frame (STEVOR) and surface-associated interspersed protein (SURFIN) families [2]. Some PfEMP1 variants cause rosetting [3–7], where the infected erythrocyte binds to uninfected erythrocytes, and there is also some limited evidence for the involvement of RIFIN and STEVOR in rosetting interactions [8,9]. Rosetting enhances microvascular obstruction [10], and has been consistently associated with severe malaria in African children [11–14]. For this reason, rosetting is considered a parasite virulence phenotype.

It has been hypothesised that antibodies which target VSAs (PfEMP1 in particular), are clinically protective [15,16]. In addition, the diversity of PfEMP1 variants, and the fact that each *P. falciparum* isolate encodes roughly 60 distinct variants, with minimal overlap between the PfEMP1 repertoires of different isolates [17], strongly suggests that PfEMP1 is an important immune target. As protection against severe malaria is acquired after just a few infections [18], naturally acquired strain-transcending antibodies against VSAs must exist if humoral responses to polymorphic parasite antigen families contribute to protection against severe malaria. Such strain-transcending antibodies, defined here as antibodies against the surface of live infected erythrocytes that recognise multiple genetically distinct parasite strains, are thus targeting conserved rather than variant-specific epitopes. Previous work shows that sera from semi-immune adults living in malaria endemic regions have broad reactivity with the infected erythrocyte surface of diverse parasite strains, as demonstrated by agglutination assays or flow cytometry [15,16,19,20]. This broad serological recognition could be due to the presence of strain-transcending antibodies, or could be due to acquisition of an extensive pool of variant-specific antibodies, and most serological studies do not differentiate between these two possibilities. A small number of studies have identified human strain-transcending monoclonal and polyclonal antibodies to VAR2CSA, the PfEMP1 variant that causes infected erythrocyte adhesion to placental syncytiotrophoblasts [21–23], and broadly strain-transcending antibodies to RIFIN variants have been described [24]. The only study that has sought to identify human strain-transcending antibodies to rosetting parasites found only variant-specific responses [6]. However, only three rosetting strains were studied [6], none of which had the dual rosetting and IgM Fc-binding adhesion phenotype that is most strongly associated with severe malaria [25–27], and which can be targeted by strain-transcending antibodies against PfEMP1 raised in rabbits [7].

We hypothesised that people living in malaria-endemic regions acquire strain-transcending IgG antibodies against virulence-associated VSAs, and sought to test this hypothesis by first identifying plasma samples from malaria-exposed humans that recognise multiple parasite strains displaying the rosetting IgM Fc-binding phenotype, and then using the broadly-reactive plasma in IgG elution experiments to test for homologous and heterologous infected erythrocyte recognition.

## Materials and Methods

### Human plasma samples and research ethics

80 anonymised archived human plasma samples from residents of malaria-endemic regions (40 adults and 40 children) were used. The samples were collected after informed consent from donors or their parents/guardians, as part of previous studies in Mali [28], Malawi [29], Papua New Guinea [30], Kenya [31], and Ghana (Y. Azasi, G. Awandare and J.A. Rowe, unpublished). The number of samples studied from each location is shown in Table 1. The children had clinical *P. falciparum* infection at the time of plasma collection, whereas the adults were asymptomatic. The negative control human serum pool was from 15 European donors at the Scottish National Blood Transfusion Service (SNBTS), Edinburgh, UK. Ethical approval was granted as described [28–31] and from the University of Edinburgh, School of Biological Sciences Ethical Review Panel (arowe002), Scottish National Blood Transfusion Service (SNBTS) Ethics Review Board (19∼6), Ghana Health Service Review Committee ID No. GHS-ERC: 03/05/14 and Noguchi Memorial Institute for Medical Research-IRB study# 020/13-14.

**Table 1.**
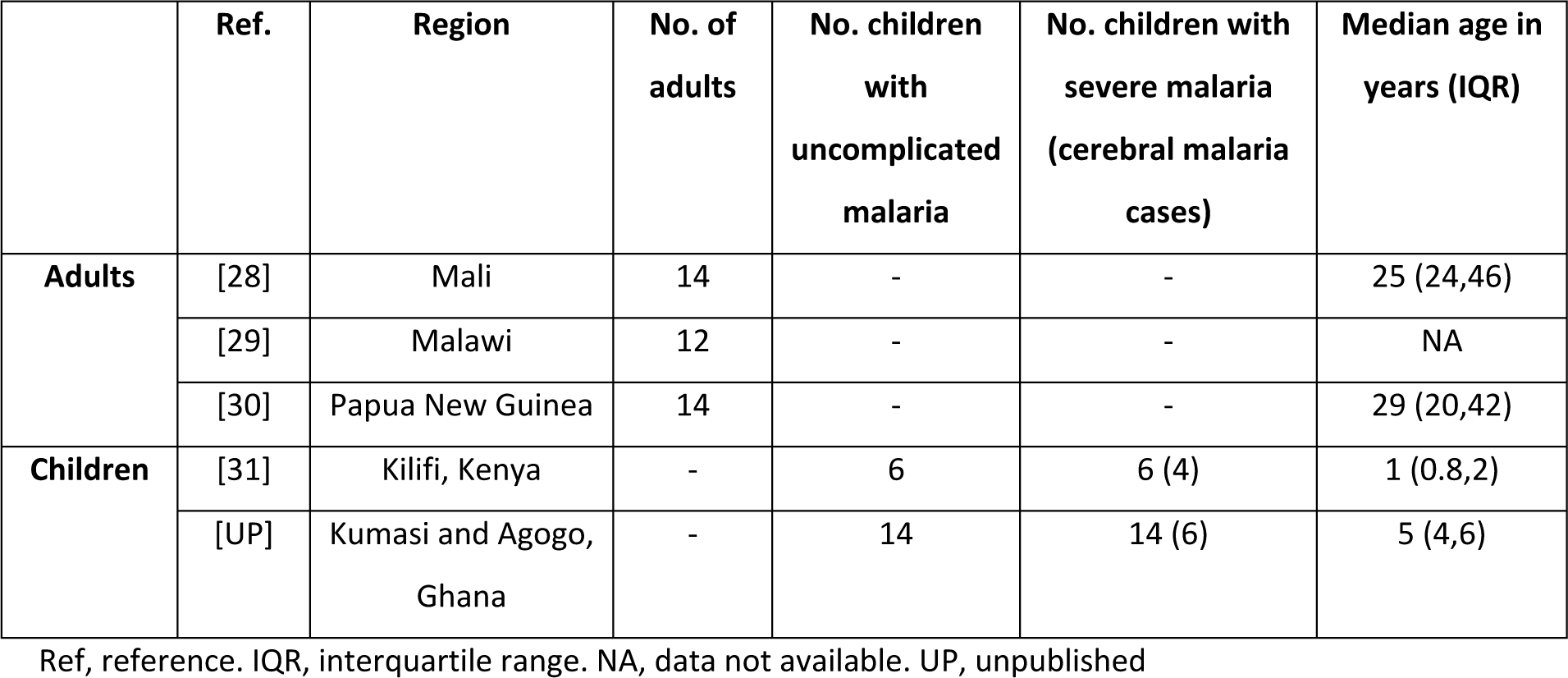
Geographical origin of human plasma samples.

|  | Ref. | Region | No. of adults | No. children with uncomplicated malaria | No. children with severe malaria (cerebral malaria cases) | Median age in years (IQR) |
| --- | --- | --- | --- | --- | --- | --- |
| **Adults** | [28] | Mali | 14 | - | - | 25 (24,46) |
|  | [29] | Malawi | 12 | - | - | NA |
|  | [30] | Papua New Guinea | 14 | - | - | 29 (20,42) |
| **Children** | [31] | Kilifi, Kenya | - | 6 | 6 (4) | 1 (0.8,2) |
|  | [UP] | Kumasi and Agogo, Ghana | - | 14 | 14 (6) | 5 (4,6) |

Ref, reference. IQR, interquartile range. NA, data not available. UP, unpublished

### Rabbit antibodies to PfEMP1

The antibodies to specific PfEMP1 variants were generated previously by immunising rabbits with recombinant DBLα protein produced in *E. coli*, then purifying total IgG from the rabbit serum on a protein A column [7][32](J.A. Rowe, unpublished data).

### P. falciparum strains

The geographical origin, *var* gene/PfEMP1 expression and adhesion phenotype of the culture-adapted parasite strains studied are shown in Table 2. In addition to the well-established laboratory strains IT, HB3, 3D7 and TM284, three recent culture-adapted Kenyan strains [24] were used. One of the Kenyan strains (11019R+) spontaneously showed high levels of rosetting, and the other two (9197 and 9605) were selected for rosetting using Percoll or gelatin sedimentation [33]. The strain 9197R+ showed two distinct phenotypic forms in parallel selections, 9197R+ which expresses the rosette-mediating PfEMP1 variant 9197varR1 (encoded by *PC0053-C.g687* in the Pf3k database [34]) (J.A. Rowe, unpublished data) and 9197varUNR+ which does not express 9197varR1, and whose predominant PfEMP1 type has not yet been identified.

**Table 2.** Geographical origin and adhesion phenotype of *P. falciparum* strains studied.

| Genotype<br>(alternative name)<br>[reference] | Origin | Parasite strain<br>(alternative names)<br>[reference] | Var gene<br>transcribed/DC*<br>(alternative name)<br>[reference] | Adhesion<br>phenotype<sup>§</sup> |
| --- | --- | --- | --- | --- |
| IT (IT4, IT04,<br>FCR3)[35] | South-East Asia<br>[36] | ITvar60R+ (R+PA1,<br>PAR+, FCR3S1)<br>[25,37] | *ITvar60*/DC11<br>(*FCR3S1.2var2*) [5,7] | IgM-positive<br>rosetting [25,26] |
|  |  | IT/R29 (R29,<br>R29R+)[38] | *ITvar9*/DC16<br>(*R29var1*) [3] | IgM-negative<br>rosetting [26] |
|  |  | IT-HBEC [39] | *ITvar19*/DC8 [32] | HBEC<sup>#</sup>-binding [32] |
| TM284 [25] | Thailand | TM284R+ [25] | *TM284var1*/DC11<br>[7,27] | IgM-positive<br>rosetting [25,26] |
| HB3 [40,41] | Honduras | HB3R+[7] | *HB3var6*/DC11 [7] | IgM-positive<br>rosetting [7] |
| 11019 (PFKE10)<br>[24] | Kenya | 11019R+ | *11019varR1*/DC11-like (*PX0203.g54*)[UP] | IgM-positive<br>rosetting [UP] |
| 9197 (PC0053-C)<br>[UP] | Kenya | 9197R+ | *PC0053-C.g687* [UP] | IgM-negative<br>rosetting [UP] |
|  |  | 9197varUNR+ | Unknown |  |
| 9605 (PFKE08) [24] | Kenya | 9605R+ | Unknown | Rosetting/IgM<br>inconclusive [UP] |
| NF54 [42] | Africa [36] | 3D7-HBEC [39] | *3D7_PFD0020c* [32] | HBEC<sup>#</sup>-binding [32] |

\*DC: domain cassette [17]. The PfEMP1 architecture and amino acid sequence identities between domains for these variants are given in references [7] and [32].
<sup>§</sup>parasite strains that have the dual rosetting and IgM Fc-binding phenotype are here designated “IgM-positive rosetting” and those that form rosettes but do not bind IgM-Fc are designated “IgM-negative rosetting”.
<sup>#</sup>HBEC: human brain endothelial cell
UP: Unpublished.

### P. falciparum culture

*P. falciparum* strains were cultured at 2% haematocrit in O+ erythrocytes and RPMI-1640 medium (Lonza, BE12-167F or Gibco, 31870-025) supplemented to give the following final concentrations: L-glutamine 2 mM (Gibco, 2530-081), glucose 16 mM, 4-(2-hydroxyethyl)-1-piperazineethanesulfonic acid (HEPES) 25 mM (Lonza, 17-737F), gentamicin 25 μg/ml (Lonza, 17-518L), AlbuMAX II Lipid-rich bovine serum albumin (BSA) (Gibco 11021037) 0.25%, and pooled human serum 5%, and the pH adjusted to 7.2-7.4 with sodium hydroxide. Both human serum and blood for culture were obtained from SNBTS, Edinburgh, UK. Cultures were gassed using 94% nitrogen, 1% oxygen and 5% carbon dioxide (BOC, 280648-L) and incubated at 37°C. Cultures were maintained at ∼2% haematocrit and 2-10% total parasitaemia. Rosetting was maintained by selection with plasmagel flotation or Percoll selection approximately once a week [33].

### Immunofluorescent staining of live infected erythrocytes with human plasma

Human plasma samples were tested against four IgM-binding rosetting parasite strains (ITvar60R+, TM284R+, HB3R+ and 11019R+)[7] and two DC8-expressing parasite strains that were previously selected for binding to human brain endothelial cells (ITvar19 and 3D7_PFD0020c)[32]. Parasite culture was washed in phosphate buffered saline (PBS)/1% BSA pre-warmed to 37°C, and then re-suspended in PBS/1% BSA at 1% haematocrit. 20 µl of washed culture suspension per well was aliquoted into a round bottomed 96 well plate (Corning, 3799), with samples assayed in duplicate. Each parasite line was tested in two independent experiments. 2.5 μl of human plasma from malaria-endemic region donors (or non-endemic European serum pool as a negative control), and 0.5 μl of 1 mg/ml rabbit IgG raised against the DBLα region of the rosette-mediating PfEMP1 variant for each parasite line [7](or non-immunised rabbit IgG as a negative control), was added and the volume in each well made up to 25 µl with PBS/1% BSA. The plates were vortexed to mix the samples and then incubated for 45 min at 37°C with re-suspension by vortexing of the plate every 15 min. The cells were washed three times with 200 µl PBS and resuspended in 40 µl of PBS/1% BSA containing 1/2500 of Vybrant^TM^ DyeCycle^TM^ Violet stain (Invitrogen, V35003) to stain parasite DNA, 1/1000 Alexa Fluor^TM^ 647-conjugated cross-absorbed goat anti-rabbit IgG (Invitrogen, A-21244) and 1/1000 Alexa Fluor^TM^ 488-conjugated cross-absorbed goat anti-human IgG (heavy chain specific) (Biotium, 20444). The plates were incubated for 30 min at 37°C in the dark, with resuspension after 15 min by gentle vortexing. The cells were washed twice in 200 µl PBS and once in PBS/1% BSA. After the final wash, the cells pellets were fixed in 50 µl of 0.5% paraformaldehyde (Thermo Fisher, 28906) in PBS for 15 min at room temperature. The paraformaldehyde was removed and the cell pellets resuspended in 200 µl of PBS containing 0.1% BSA, 0.01% sodium azide, and 100 µg/ml fucoidan (Molekula, 9072-19-9), to disrupt rosettes [7]. The plates were protected from light and stored at 4°C until analysis by flow cytometry within 24 h of staining.

#### Data acquisition and gating strategy for human plasma experiments

The 96 well plates were read on a BD LSRFortessa^TM^ instrument (BD Biosciences) using lasers at 405 nm (for Vybrant^TM^ DyeCycle^TM^ Violet), 488 nm (for Alexa Fluor^TM^ 488) and 640 nm (for Alexa Fluor^TM^ 647). 100 μl from each well was analysed at a flow rate of 2 μl/s until a maximum of 500,000 events were recorded, or 50s had elapsed, whichever came first. For each sample, at least 400 cells from the population of interest (i.e., PfEMP1-positive infected erythrocytes) were recorded. The data were analysed in FlowJo v.10 (BD Biosciences). All events were gated using forward and side scatter to exclude debris, followed by gating on all infected erythrocytes (Vybrant^TM^ DyeCycle^TM^ Violet high), followed by PfEMP1 positive cells (Alexa Fluor^TM^ 647 high), to identify the rosetting PfEMP1 variant-expressing infected erythrocyte cell population of interest. Human IgG bound to these cells was detected in the 488 channel and the median fluorescence intensity (MFI) was recorded. An example of the gating strategy is shown in Figure 1.

**Figure 1.**
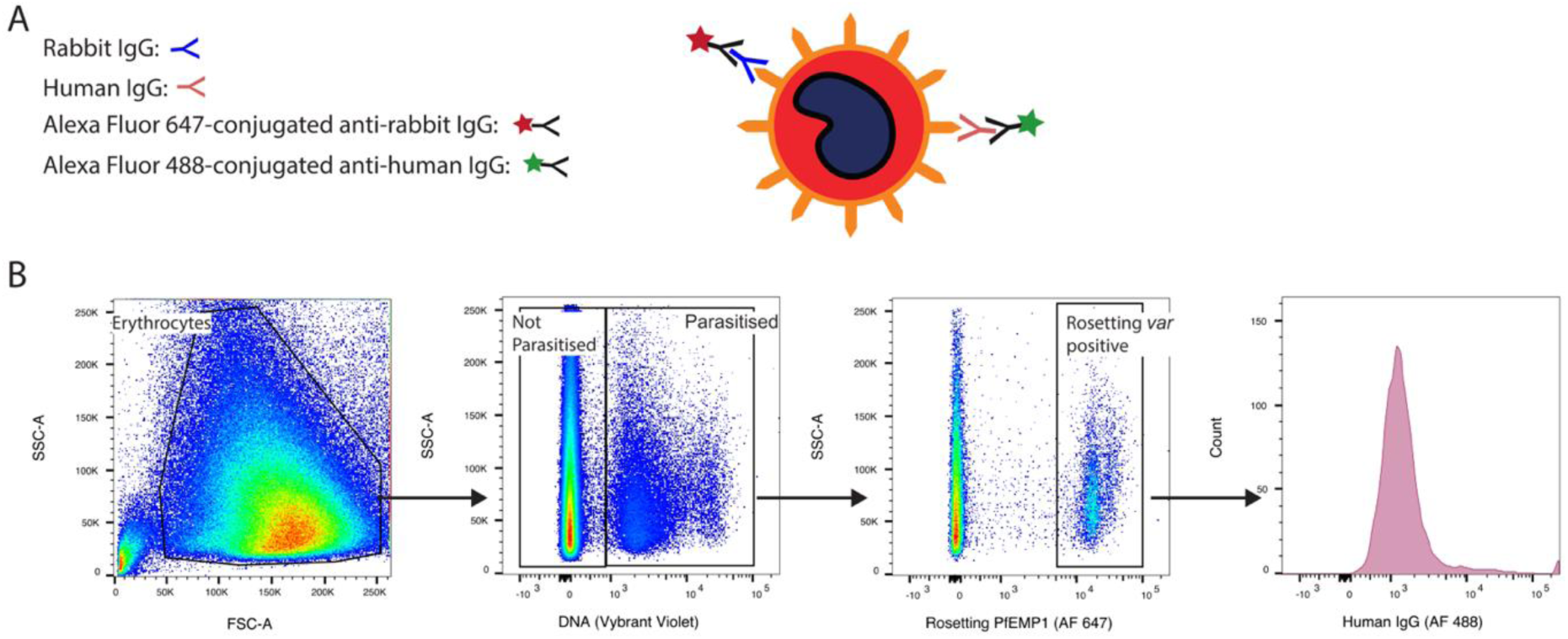
Detection of naturally acquired human IgG to rosetting parasite strains by flow cytometry. A) Schematic diagram of the staining strategy. B) Gating strategy. Forward and side scatter were used to exclude debris and gate on all erythrocytes (from left, first panel). All parasitised erythrocytes were identified by DNA staining with Vybrant^TM^ Dyecycle^TM^ Violet (second panel), and those that stained surface positive for the rosette-mediating PfEMP1 variant of interest were detected with 20 μg/ml rabbit polyclonal IgG against the NTS-DBLα domain of the PfEMP1 variant and an Alexa Fluor^TM^ 647-conjugated anti-rabbit IgG secondary antibody at 1/1000 (third panel). The median fluorescence intensity (MFI) of the Alexa Fluor^TM^ 488 channel (human IgG stain) for this “rosetting *var* positive” population was quantified (fourth panel). Human plasma was used at 1/10 dilution and Alexa Fluor^TM^ 488-conjugated anti-human IgG (gamma chain) was used at 1/1000 concentration. AF, Alexa Fluor^TM^.

#### Data analysis and visualisation for human plasma experiments

Because human plasma/sera sometimes show non-specific binding to infected and/or uninfected erythrocytes, controls were carried out and fluorescence intensity measurements were corrected for background staining (cMFI) as described [43]. The cMFI = (MFI of the PfEMP1-positive infected erythrocytes stained with endemic plasma – MFI of the uninfected erythrocytes from the same sample) - (MFI of the PfEMP1-positive infected erythrocytes stained with the European negative control serum pool – MFI of the uninfected erythrocytes from the same sample). An example is shown in Figure S1. To visualise the data, the cMFI values for all plasma samples against all parasite strains were summarised in a checkerboard. To quantify the seroprevalence (percentage of plasma samples giving a positive result) for each parasite strain, a cMFI of greater than 2 standard deviations (SD) of the MFI of the PfEMP1-positive population with the European negative control was considered to be positive. This SD of the negative control for each parasite line was calculated from four independent experiments using the mean MFI of duplicate negative control wells from each. Furthermore, all cMFI values below 100 fluorescence units (fu) were considered negative as recognition was not compelling for samples when the cMFI was <100, even if this was above two standard deviations of the negative control (Figure S2). This additional cut-off was applied to reduce the incidence of type 1 error. To examine associations between cMFI and disease severity, any negative cMFI values were converted to zero, then Mann Whitney U tests were performed in GraphPad Prism v8 (GraphPad Software), without correction for multiple comparisons. To compare the number of positive samples in uncomplicated versus severe malaria, Fisher’s exact test was performed using GraphPad Prism v8.

### Magnetic-activated cell sorting (MACS) to purify infected erythrocytes

Infected erythrocytes of strains ITvar60R+ or TM284R+ were purified by MACS prior to adsorption with human plasma. RPMI 1640 containing 2 mM L-glutamine and 25mM HEPES, but lacking bicarbonate (Gibco, 13018-015) was supplemented to 16 mM glucose and 25 μg/ml gentamicin and adjusted to pH 7.2-7.4 with sodium hydroxide to make incomplete binding medium (ICBM). 0.5% immunoglobulin-free bovine serum albumin (BSA) (Sigma-Aldrich, A0336) and either 1 mg/ml heparin (Sigma-Aldrich, H4784-1G) for ITvar60R+ or 50 μg/ml fucoidan (Molekula, 9072-19-9) for TM284R+, was added to ICBM to make rosette-disrupting medium (RDM). CS columns (Miltenyi Biotec, 120-000-490) were fitted with a three-way tap with a 20G needle attached to the outflow port and the column applied to a VarioMACS^TM^ Separator (Miltenyi Biotec, 130-090-282). The column was primed with 10 ml PBS, then 20 ml ICBM, followed by 5 ml rosette-disrupting medium. 400-500 μl packed cell volume of parasite culture at 5-10% parasitaemia was washed in ICBM and resuspended in 10 ml rosette-disrupting medium before being applied to the column and run through at ∼1 drop per second. The flow through was collected and run through the column a second time. The column was washed with 20 ml of rosette-disrupting medium, removed from the VarioMACS^TM^ Separator, and 10 ml of rosette-disrupting medium used to elute the adherent infected erythrocytes. 20-50 μl of rosetting infected erythrocytes were obtained, with a parasitaemia of 60-70% achieved for most purifications (>40% required for use in experiments). Eluted infected erythrocytes were washed twice in 20 ml of ICBM to remove heparin or fucoidan then resuspended in ICBM.

### Adsorption and antibody elution from the infected erythrocyte surface

Eight individual plasma samples (M2, M3, M4, M6, G7, G9, G12, G14) showing broad recognition of rosetting strains were used in adsorption and antibody elution experiments. Prior to adsorption, any antibodies recognising uninfected erythrocytes in each plasma sample were pre-adsorbed by incubation with an equal packed cell volume of O+ erythrocytes from a Scottish donor at 20 rpm on a rotating wheel for 1 hour at room temperature, and the process repeated with fresh erythrocytes a total of 4 times. The pre-adsorbed plasma was stored at 4°C until use. The antibody adsorption and elution method was based on the procedure previously described by Moll *et al*. [44]. 20-50 μl PCV of MACS purified infected erythrocytes of strains ITvar60R+ or TM284R+, were resuspended in 100 μl of ice-cold pre-absorbed plasma and incubated on ice for 90 min with gentle re-suspension every 10 min. The cells were pelleted by centrifugation at 9447 xg for 5 s and the supernatant removed. The cells were washed five times by resuspending in 1 ml of ice-cold 0.15 M NaCl by gentle pipetting, re-pelleting by centrifugation as above, and removal of the wash buffer. After the fifth wash, cell pellets were resuspended in 50 μl ice-cold 0.15 M NaCl. Three rounds of elution were undertaken at successively lower pH points. 0.2 M HCl was added to 0.2 M glycine/0.2 M NaCl until solutions at the desired pH were obtained (pH 2.7, 2.2 and 1.7). First, 50 μl ice-cold glycine-HCl at pH 2.7 was added to the resuspended cells and the suspension mixed gently by gentle pipetting before being placed on ice for 2 min. The cells were pelleted by centrifugation as described above, and the supernatant neutralised by transferring to a fresh tube containing 3 μl 2M ice-cold Tris-base. After re-suspension of the cell pellet in 50 μl ice-cold 0.15M NaCl, the elution was repeated with 50μl glycine-HCl at pH 2.2, with the supernatant transferred into a tube containing 6μl 2M Tris-base. This was then repeated once more with glycine-HCl at pH 1.7 and transfer of the supernatant into a tube containing 8 μl 2M Tris-base. The pH of the eluted fractions was checked using pH strips and adjusted to pH 7 by the addition of further 2M Tris-base as required. All three eluted fractions were centrifuged at 9447 xg for 5 min to pellet debris and the three fractions pooled into a clean tube. The eluted antibody was buffer-exchanged into PBS using a Millipore Amicon ultra 100K microcentrifuge concentration unit (Merck, UFC510024) with four rounds of buffer-exchange using 450 μl of PBS per round. Purified eluate was recovered in roughly 20 μl total volume by inverting the Millipore Amicon ultra 100K microcentrifuge concentration unit in a clean microcentrifuge tube and spinning for 2 min at 1000 x g. The A280 protein concentration in the purified eluate was measured using a Nanodrop Spectrophotometer ND-1000 and recorded before the addition of 1 μl of BSA 30% to give a final concentration of roughly 1% BSA. The purified eluate was stored at 4°C until use. For most samples, eluate volume was adjusted with PBS to 150μl total volume to give sufficient volume for downstream experiments, resulting in protein concentrations up to 46 μg/ml. The exceptions were M2 eluates from both ITvar60R+ and TM284R+, which were adjusted to total volumes of 216 μl and 180 μl respectively. This was to bring their protein concentrations to below 200 μg/ml, which was the concentration used for the human IgG negative control (Sigma, I2511) in subsequent experiments. These diluted eluates were used neat in the staining experiments.

### Immunofluorescent staining of infected erythrocytes with eluted antibodies

Eight rosetting parasites strains (Table 2) were stained with eluted antibodies. Staining for recognition of the surface of live infected erythrocytes was carried out as described above for staining with human plasma, with modifications as follows: i) 10 μl of washed culture suspension at 1% haematocrit was aliquoted into each well of the 96 well plate, the PBS/1% BSA removed, and 12.5 μl of the neat eluate or negative control commercially available purified human IgG (Sigma, I2511) in PBS at 200 μg/ml was used to resuspend the cells. ii) the samples were not assayed in duplicate due to limited eluate availability and each parasite strain was tested once. iii) the staining strategy did not include variant specific rabbit polyclonal IgG. Instead, the secondary antibody 1/1000 Alexa Fluor 647^TM^-conjugated goat anti-human IgG (gamma chain specific) (Biotium, 20448) in PBS/1% BSA contained 1/2500 of Vybrant^TM^ DyeCycle^TM^ Violet stain and 20 μg/ml Ethidium Bromide (Sigma, 46067). The inclusion of both Vybrant^TM^ DyeCycle^TM^ Violet (DNA stain) and Ethidium Bromide (DNA and RNA stain) allows for gating to distinguish mature infected erythrocytes (pigmented trophozoites and schizonts) (DNA and RNA high) from uninfected erythrocytes (DNA and RNA low) and ring-stage infected erythrocytes (DNA high, RNA low), as previously described [45].

#### Data acquisition and gating strategy for eluted antibody experiments

Data acquisition was as described above for human plasma samples, in this case using lasers 405 nm (Vybrant^TM^ DyeCycle^TM^ Violet), 488nm (Ethidium bromide) and 605 nm (Alexa Fluor 647^TM^). All events were gated using forward and side scatter to exclude debris, followed by gating on mature infected erythrocytes (Vybrant^TM^ DyeCycle^TM^ Violet high, Ethidium bromide high) to identify the mature-infected erythrocyte cell population of interest. Human IgG bound to these cells was detected in the 647 channel and the median fluorescence intensity was recorded.

#### Data analysis and visualisation for eluted antibody experiments

For the eluted antibody experiments, cMFI = (MFI of the mature infected erythrocytes stained with eluted antibody - MFI of the uninfected erythrocytes from the same sample) - (MFI of mature infected erythrocytes stained with the commercial negative control human IgG – MFI of the uninfected erythrocytes from the same sample). To visualise the data, the cMFI values for each eluate against each parasite strain were summarised in a checkerboard.

### Rosette disruption with eluates

A 200μl aliquot of parasite culture at ∼2% Ht was stained with 25 μg/ml Ethidium Bromide for 5 min at 37°C. The supernatant was removed and the cells resuspended at 4-5% haematocrit in RPMI incomplete binding medium (as described above for magnetic-activated cell sorting) supplemented to 15% heat-inactivated normal human serum from Scottish donors. 2μl of PBS or eluate were added to 2μl aliquots of the pre-stained culture suspension and incubated for 30 min at 37°C with resuspension at 15 min. The negative controls were the “no antibody” control (with PBS added) and the “eluate negative” sample M4 (IT), as this eluate did not recognise any heterologous parasites by flow cytometry, so would not be expected to disrupt rosettes in heterologous strains. As a positive control, 2 μl of 1mg/ml rabbit IgG against the specific PfEMP1 variant for each parasite line [7] was added to an aliquot of culture. For 9197varUNR+, antibodies are not available, so heparin (Sigma-Aldrich, H4784-1G) at a final concentration of 1mg/ml was used as a positive control. The aliquot identities were masked before counting to prevent observer bias. Wet preparations were made on multi-spot slides and at least 200 mature infected erythrocytes per spot were observed using a Leica DM2000 fluorescence microscope and 40x objective, with simultaneous brightfield/fluorescence to visualise both infected and uninfected erythrocytes. The rosette frequency (percentage of mature infected erythrocytes binding two or more uninfected erythrocytes) was recorded. Three independent experiments were done for each parasite strain/eluate combination as follow: TM284R+ with eluate M2(IT); 9197varUNR+ with eluate M2 (IT), M3 (IT), M6 (IT) and M2(TM284); HB3R+ with eluate M3 (IT). Not all parasite strain/eluate combinations could be tested due to limited eluate availability. The mean rosette frequency in the presence of eluted antibody was compared to that in the M4 “eluate negative” control using Student’s t test or one-way ANOVA in GraphPad Prism v8.

### Data availability

All raw data are available from the authors on request.

## Results

### Detection of naturally acquired human IgG to rosetting parasite strains

First, we investigated whether individuals living in malaria endemic regions naturally acquire antibodies to multiple rosetting parasite strains. Archived plasma samples from 40 adults and 40 children (Table 1) were tested for IgG recognition of the surface of live infected erythrocytes by flow cytometry, using four rosetting IgM Fc-binding parasite strains (Table 2). Because of spontaneous *var* gene switching [38,46], cultured parasites contain a heterogeneous mix of infected erythrocytes expressing different PfEMP1 types, even after selection. Therefore, we used a dual human plasma/rabbit anti-PfEMP1 staining protocol to allow specific examination of human IgG binding to infected erythrocytes expressing the rosette-mediating PfEMP1 types of interest. For each parasite strain, the PfEMP1-positive infected erythrocytes were identified using rabbit polyclonal IgG against the rosette-mediating PfEMP1 N-terminal domain and an Alexa Fluor^TM^ 647-conjugated anti-rabbit IgG secondary antibody. The human antibodies bound to this infected erythrocyte population were detected with an Alexa Fluor^TM^ 488-conjugated anti-human gamma chain-specific secondary antibody (Figure 1). Any background staining of infected or uninfected erythrocytes, which occurs commonly with human sera and plasma [43], was corrected for as described in the methods and Figure S1. Examples of human plasma samples showing negative or strong positive IgG staining are shown in Figure 2.

**Figure 2.**
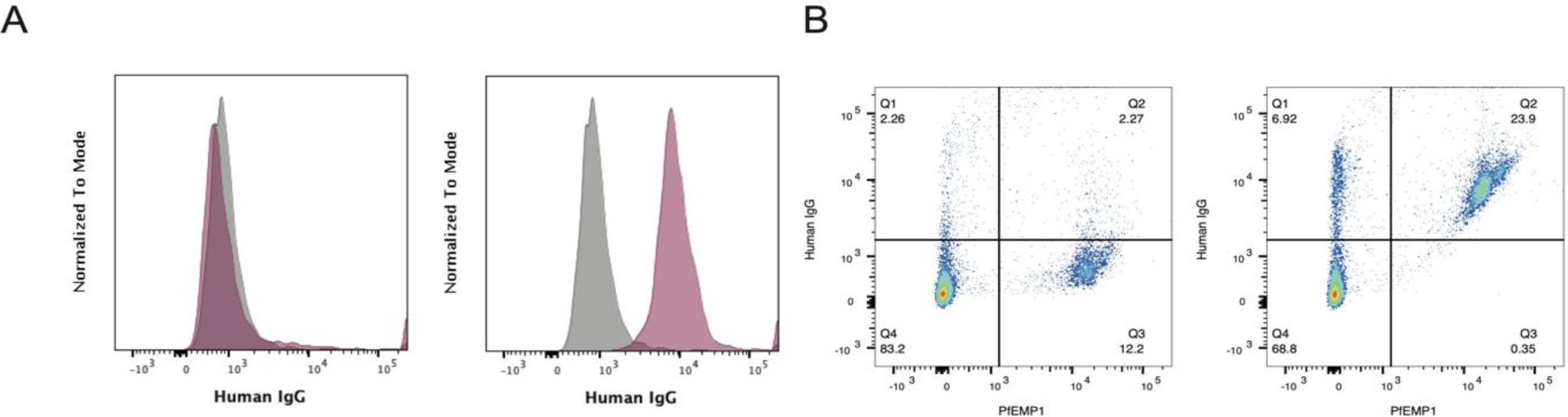
Examples of negative and positive IgG responses to infected erythrocytes. A) Fluorescence intensity histograms of rosetting PfEMP1-positive cell populations exemplifying plasma samples, red, with negative reactivity (plasma MW11 to parasite 11019R+, left), and strong positive reactivity (plasma M3 to parasite 11019R+, right). Background staining with the negative control European serum pool is shown in grey. B) Dot-plots of samples shown in A but rather than only rosetting PfEMP1 positive cells, all infected erythrocytes are shown. Human plasma and sera were used at 1/10 dilution. An Alexa FluorTM 488-conjugated anti-human IgG (gamma chain) antibody was used at a 1/1000 dilution to detect human IgG. Surface staining for PfEMP1 was with 20 μg/ml rabbit polyclonal IgG raised against the NTS-DBLα domain of the PfEMP1 variant of interest and detected with an Alexa FluorTM 647-conjugated anti-rabbit IgG secondary antibody at 1/1000 dilution.

The four rosetting parasite strains, which originated from SE Asia, Thailand, Honduras and Kenya, were well-recognised by adult plasma samples from Mali, Papua New Guinea and Malawi (Figure 3). Seroprevalence values, defined here as the percentage of plasma samples positive for recognition of each strain, were calculated and 93% (37/40) of the adult samples recognised at least one rosetting strain, with 58% (23/40) recognising all four rosetting strains (Table 3). For the children’s plasma samples, which came from donors aged 4 months to 11 years, 40% (16/40) recognised at least one rosetting strain, but only 1/40 (3%) recognised all four rosetting strains (Figure 3 and Table 3). For comparison, IgG responses to two *P. falciparum* strains expressing virulence-associated DC8 variants 3D7_*PFD0020c* and *ITvar19* [32], were tested for recognition with the same plasma set (Figure 4 and Table 3). 85% (34/40) of adult samples and 53% (21/40) of children’s samples recognised at least one of the DC8-containing parasite strains, and 63% (25/40) of adults and 30% (12/40) of children recognised both strains.

**Figure 3.**
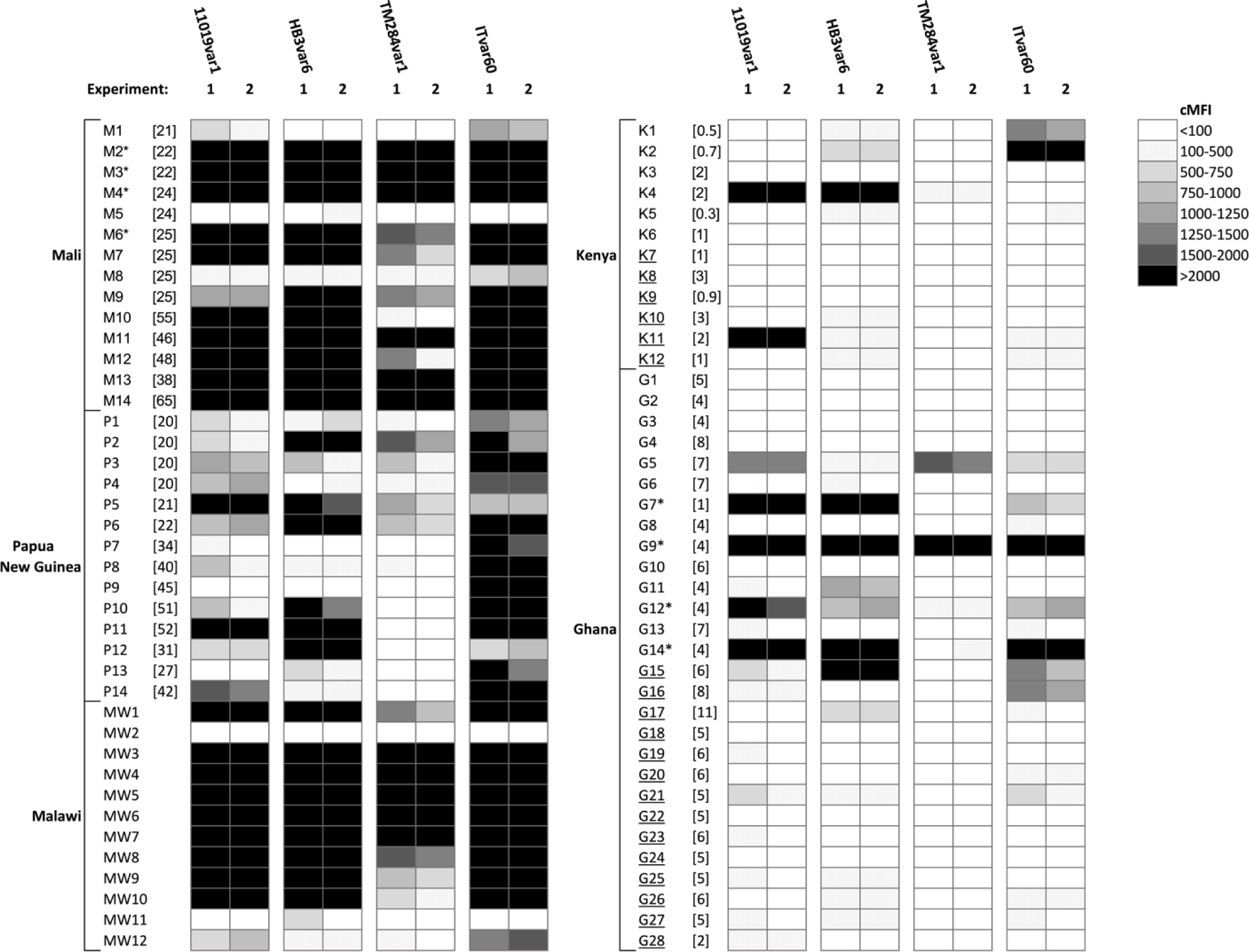
Recognition of infected erythrocytes expressing rosetting PfEMP1 variants by human IgG. Corrected median fluorescence intensities (cMFI) of rosetting PfEMP1 variant-expressing infected erythrocytes, incubated with human plasma at 1/10 dilution, with bound IgG detected with an Alexa Fluor^TM^ 488-conjugated anti-human IgG (gamma chain) antibody at 1/1000 dilution. Left panel, adults’ plasma, right panel, childrens’ plasma. The age in years of each individual at the time of plasma donation is shown in square brackets, if known. Each plasma sample was tested in duplicate in each experiment and the mean of the duplicates is presented. The results of two independent experiments are shown. Samples from children with severe malaria are underlined. *Samples used in eluted antibody experiments.

**Figure 4.**
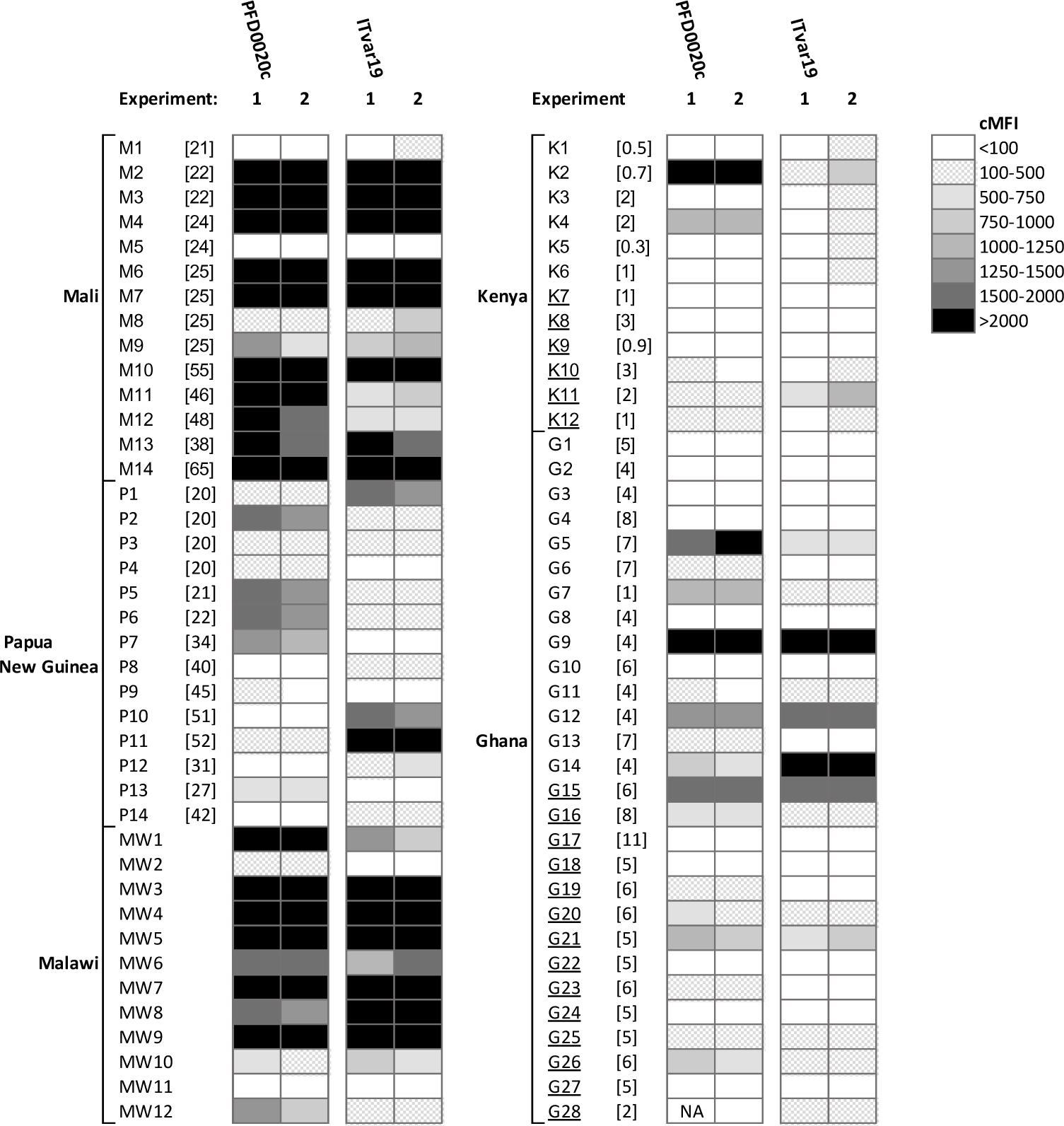
Recognition of infected erythrocytes expressing DC8 PfEMP1 variants by human IgG. Corrected median fluorescence intensities (cMFI) of DC8 PfEMP1 variant expressing infected erythrocytes, incubated with human plasma at 1/10 dilution, with bound IgG detected with an Alexa Fluor^TM^ 488-conjugated anti-human IgG (gamma chain) antibody at 1/1000 dilution. Left panel, adults’ plasma, right panel, childrens’ plasma. The age in years of each individual at the time of plasma donation is shown in square brackets, if known. Each plasma sample was tested in duplicate in each experiment and the mean of the duplicates is presented. The results of two independent experiments are shown. NA, data not available due to technical error. Samples from children with severe malaria are underlined.

**Table 3.** Seroprevalence* against rosetting and DC8-expressing parasite strains.

|  | 11019varR1 | HB3var6 | TM284var1 | ITvar60 | PFD0020c | ITvar19 |
| --- | --- | --- | --- | --- | --- | --- |
| **Adults** | 83% (33) | 80% (32) | 58% (23) | 93% (37) | 70% (28) | 80% (32) |
| **Children** | 23% (9) | 30% (12) | 5% (2) | 25% (10) | 38% (15) | 50% (20) |

\*Seroprevalence is the percentage of plasma samples positive for recognition of each strain. The number of positive samples out of the 40 tested is given in parentheses. Samples were considered positive when the mean corrected MFI (cMFI) from the two staining experiments was >100 fluorescence units and was greater than two standard deviations of the European non-immune negative control values for that strain.

### IgG responses to rosetting strains and malaria severity in children

Although this was not the primary aim of the work, and the study was underpowered to detect statistically significant differences, we examined the human plasma staining data for any trends in the relationship between disease severity and the presence of IgG responses to the surface of infected erythrocytes. Previous studies suggest that children with severe malaria have lower malaria-specific immunity and are less likely to have antibodies to commonly-occurring VSAs than children with uncomplicated disease [47–50]. Of the 40 plasma samples from children, 20 were from children with uncomplicated malaria and 20 were from children with severe malaria. There were no statistically significant differences in cMFI between children with severe and uncomplicated malaria for any of the parasite strains tested (Figure S3). The rosetting strain ITvar60R+ showed the biggest difference in median values, with higher recognition by the uncomplicated malaria samples, a potentially interesting finding that could be examined further in a larger sample set. When the mean cMFI of all rosetting parasite strains for each child was used as a composite measure of recognition, there was still no statistically significant difference between children with severe and uncomplicated malaria (Figure S3: All rosetting strains). Although more children with uncomplicated malaria had a positive response to at least one of the rosetting strains than those with severe malaria (10/20 vs 4/20 children respectively), this difference was not statistically significant (Fisher’s exact test p = 0.096). There was also no statistically significant difference in cMFI between children with severe and uncomplicated malaria for the two strains expressing DC8-like PfEMP1 variants (Figure S3). Associations with distinct severe malaria syndromes such as cerebral malaria were not examined due to the small sample size.

### Detection of strain-transcending activity against rosetting strains using eluted antibodies

Plasma from four adults (M2, M3, M4 and M6) and four children (G7, G9, G12, and G14) were chosen for eluted antibody experiments, on the basis of broad recognition (positive with 3 or 4 rosetting strains) and plasma availability. Each individual plasma sample was incubated with purified infected erythrocytes and the bound immunoglobulin was eluted from the infected erythrocyte surface using cold acid-elution [19,44]. ITvar60R+ and TM284R+ were selected as the adsorbing parasite strains due to the relative ease of growing large culture volumes and maintaining the rosetting phenotype in these strains. The eluates were tested for their ability to stain homologous and heterologous live infected erythrocytes as shown in Figure 5A. As the IgG in the eluates was present at very low concentrations, co-staining with rabbit IgG to the PfEMP1 variant of interest was not included in the staining strategy, to prevent the possibility of the rabbit IgG competing with the human IgG for PfEMP1. Instead, the mature PfEMP1-expressing infected erythrocyte population was identified by staining with both Vybrant^TM^ DyeCycle^TM^ Violet and Ethidium Bromide as previously described [45] (Figure 5B), and the fluorescence intensity values were corrected for non-specific background staining as described in the methods.

**Figure 5.**
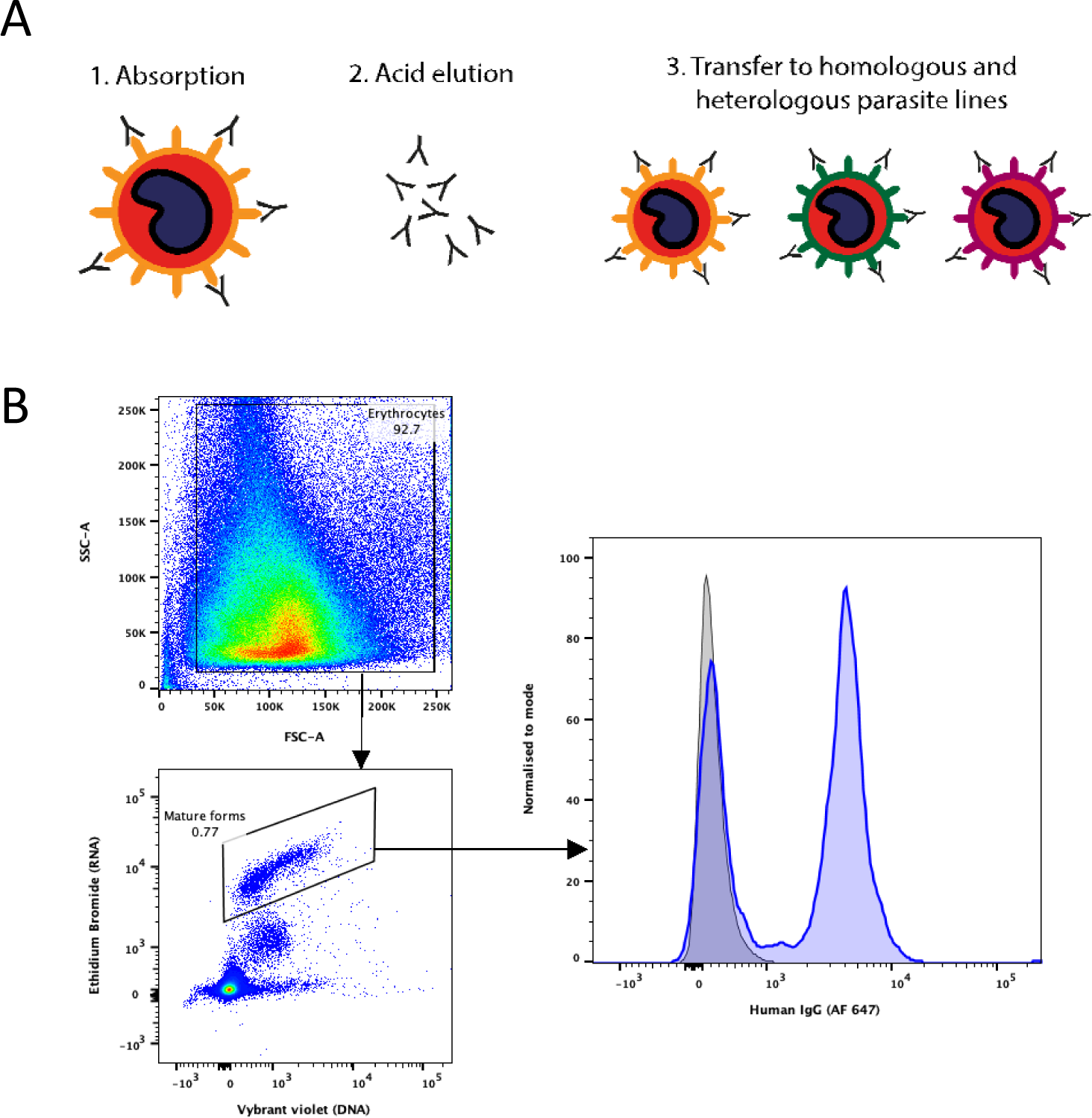
Method of detection of strain-transcending IgG in eluates. (A) Schematic diagram of the acid elution method. (B) Successful detection of IgG in eluates. Left, dot-plots showing the gating strategy used to identify mature infected erythrocytes in whole cultures. Forward and side scatter were used to exclude debris and set a gate on all erythrocytes, then mature pigmented-trophozoite and schizont infected erythrocytes were identified using Vybrant^TM^ Dyecycle^TM^ Violet (DNA stain, 1/2500 dilution) and Ethidium Bromide (DNA/RNA stain, 20 μg/ml) as the DNA and RNA high population (38). Right, fluorescence intensity histogram of ITvar60R+ mature pigmented trophozoites stained with neat eluate of plasma M2 from ITvar60R+ (blue), compared to a concentration-matched human IgG control (grey). Bound human IgG was detected with an Alexa Fluor^TM^ 647-conjugated anti-human IgG (gamma chain) antibody at 1/1000 dilution.

Using ITvar60R+ as the adsorbing strain, all eight eluates recognised the homologous parasites, showing successful elution of IgG against the infected erythrocyte surface (Figure 5B and Figure S3). For TM284R+, five eluates did not stain homologous parasites, therefore these samples were excluded, as this indicated failure to elute sufficient IgG for detection. The other three positive TM284R+ eluates and the eight positive ITvar60R+ eluates were used to stain seven geographically diverse heterologous rosetting strains (Table 2), to look for strain-transcending activity. Nine of the eleven eluates had heterologous activity against at least one other rosetting strain, with six of these reacting with multiple heterologous strains (Figure 6 and Figure 7). The strain-transcending activity of eluates from ITvar60R+ was most frequent against 11019R+, with five of the eight eluates recognising this strain. The other rosetting strains were recognised by between two and four of the ITvar60R+ eluates, except for rosetting strain IT/R29R+, which was not recognised. Given that ITvar60R+ and IT/R29R+ are different variants from the same parasite genotype (IT/FCR3), it is perhaps not surprising that they are antigenically distinct, because shared epitopes within the same repertoire would reduce the number of immunologically novel variants available to facilitate parasite survival and prolong infection in the human host. For the TM284R+ eluates, strain-transcending activity against ITvar60R+ was most frequent, as all three eluates recognised this rosetting strain. The relatively high sequence similarity between the N-terminal DBLα domain (63% amino acid identity) and Cysteine-Rich Interdomain Region (CIDRβ, 82% amino acid identity) of these two variants has been noted previously [7].

**Figure 6.**
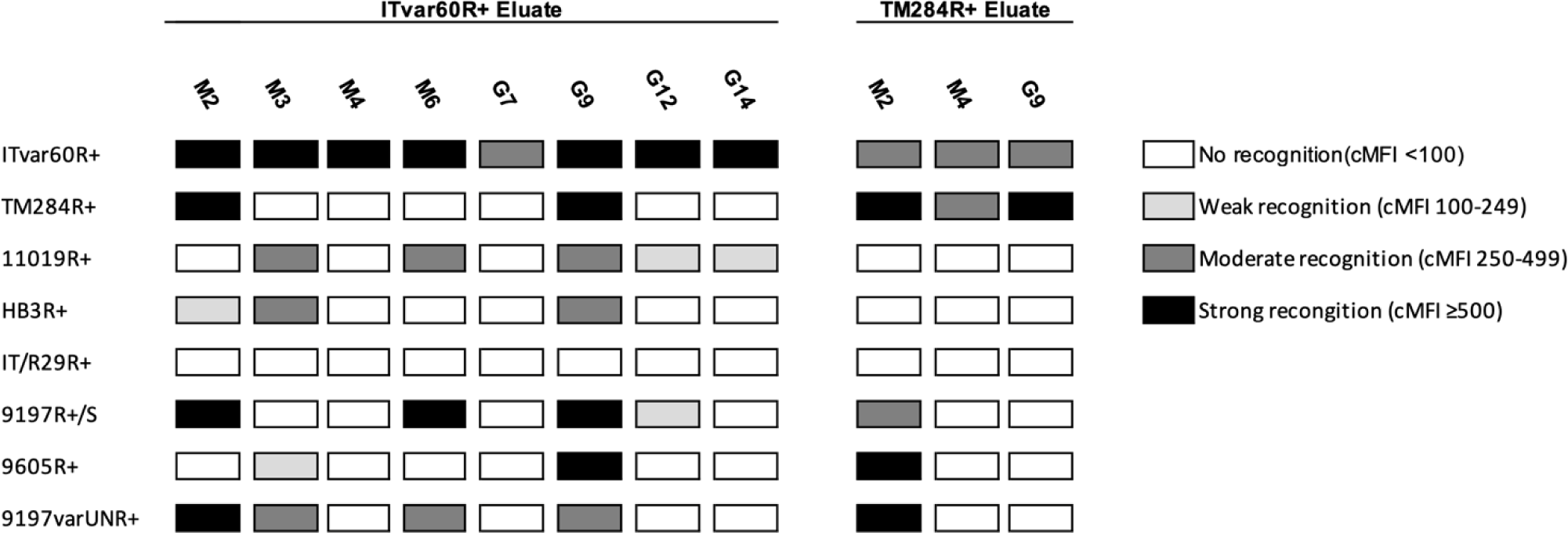
Summary of recognition of homologous and heterologous rosetting infected erythrocytes by eluted antibodies using flow cytometry. Eight rosetting parasite strains were stained with antibodies eluted from ITvar60R+ (8 individual donors; 4 adults M2, M3, M4, M6 and 4 children G7, G9, G12, G14) or from TM284R+ (3 individual donors; 2 adults M2 and M4 and 1 child G9). Mature infected erythrocytes were identified using Vybrant^TM^ Dyecycle^TM^ Violet at 1/2500 dilution (DNA stain) and Ethidium Bromide at 20μg/ml (DNA/RNA stain) (38). Eluates were used neat, and human IgG bound to mature infected erythrocytes was detected with an Alexa Fluor^TM^ 647-conjugated anti-human IgG (gamma chain) antibody at 1/1000 dilution. Corrected median fluorescence intensities (cMFI) were adjusted for background staining as described in the methods. Due to the limited availability of eluate, the experiments were performed once.

**Figure 7.**
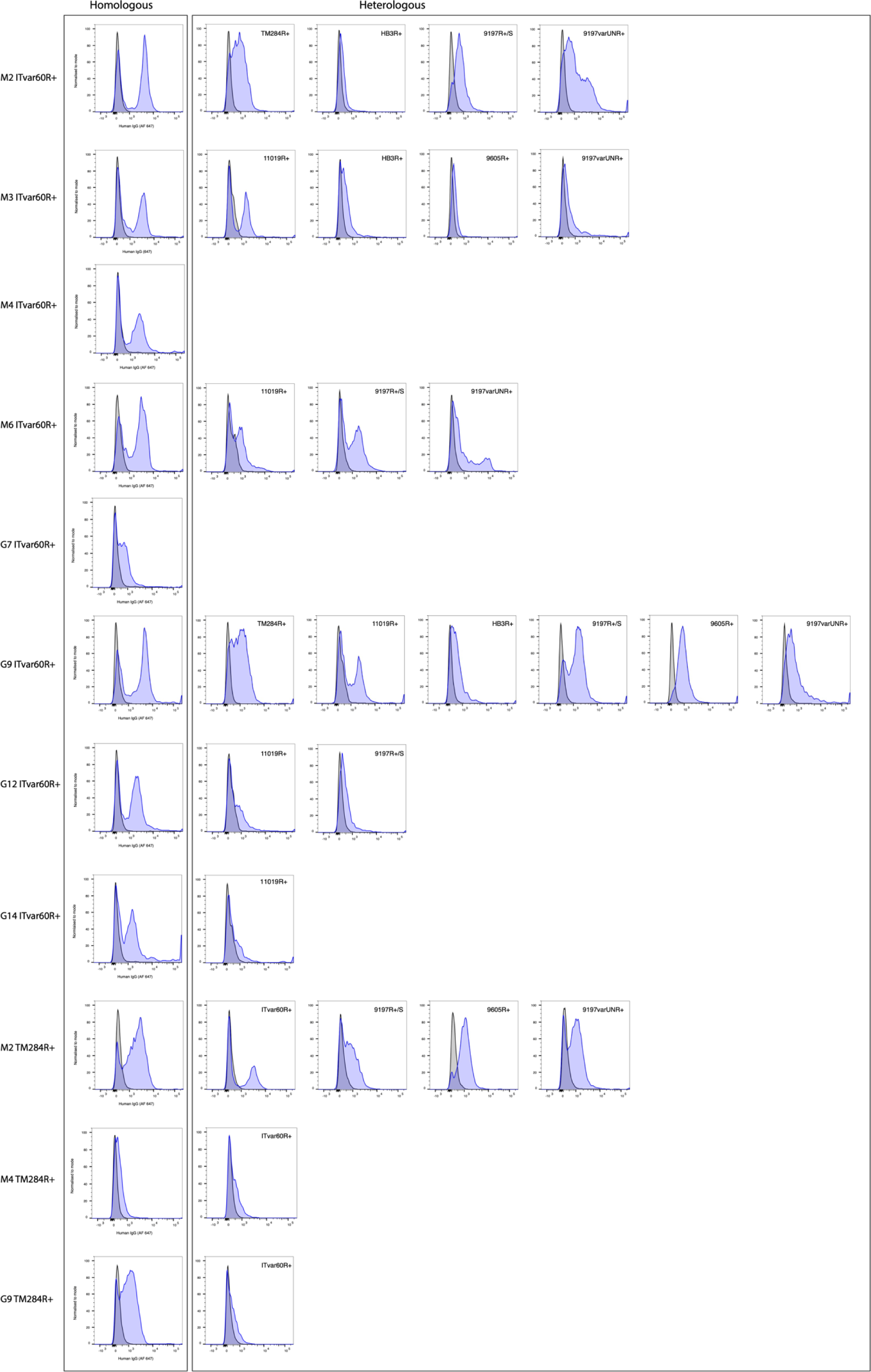
Recognition of homologous and heterologous strains by eluates. Fluorescence intensity histograms of mature infected erythrocytes, incubated with eluate (blue) or a human IgG control (grey). Each row of plots shows the results for one eluate, with the plasma and adsorbing line indicated to the left of each row. Only positive (cMFI ≥100) results are shown. Heterologous responses are displayed with the line indicated inside the histogram box. Mature infected erythrocytes were identified using Vybrant^TM^ Dyecycle^TM^ Violet at 1/2500 dilution (DNA stain) and Ethidium Bromide at 20 μg/ml (DNA/RNA stain) (38). Eluates were used neat, and human IgG bound to mature infected erythrocytes was detected with an Alexa Fluor^TM^ 647-conjugated anti-human IgG (gamma chain) antibody at 1/1000 dilution.

The pattern of staining with the eluted antibodies allowed us to draw inferences about the nature of the conserved determinant(s) on the surface of infected erythrocytes. The entire mature-infected erythrocyte population was examined, which (as described above) consists of a mixture of cells expressing the rosetting PfEMP1 variant of interest and cells that have switched to express other PfEMP1 types. If the eluted antibodies were specifically recognising the rosette-mediating PfEMP1 variant-expressing cells, then some but not all infected erythrocytes should be recognised, resulting in a bimodal distribution on the fluorescence intensity histograms. If, on the other hand, the eluted antibodies were recognising a conserved epitope or protein present on all infected erythrocytes, then a single population of positively stained infected cells would be seen. Examination of the data shows a clear bimodal pattern of IgG reactivity in many cases, as would be expected for antibodies against a VSA such as PfEMP1 (Figure 7).

### Eluate M2(ITvar60R+) can disrupt rosettes

As well as being an immune target, PfEMP1 is the adhesion molecule that mediates rosetting in many *P. falciparum* strains [3–7]. We therefore investigated whether the eluted antibodies had rosette-disrupting activity against heterologous strains, as this might be detected if they are targeting PfEMP1. Due to limited eluate availability, only some eluate/parasite line combinations were tested. Eluate M4 (eluted from ITvar60) was used as a negative control, as it only recognised the homologous line by flow cytometry, so it should not show any heterologous rosette-disrupting activity. Rabbit IgG raised to the NTS-DBLα domain of the rosette-mediating PfEMP1 variants for each strain was used as a positive control for rosette disruption, except for strain 9197varUNR+, where heparin was used. When incubated at a final dilution of 1 in 2, eluate M2 (eluted from ITvar60R+) disrupted rosettes in both TM284R+ and 9197UNR+ (Figure 8). This suggests that in the M2 plasma sample (eluted from IT parasites), PfEMP1 could be the target of the strain-transcending antibodies. The other eluates tested did not have statistically significant rosette disrupting activity on the parasite strains tested, either because they recognise epitopes or antigens distinct from the rosetting binding site, or because the concentration of IgG was too low to disrupt rosettes.

**Figure 8.**
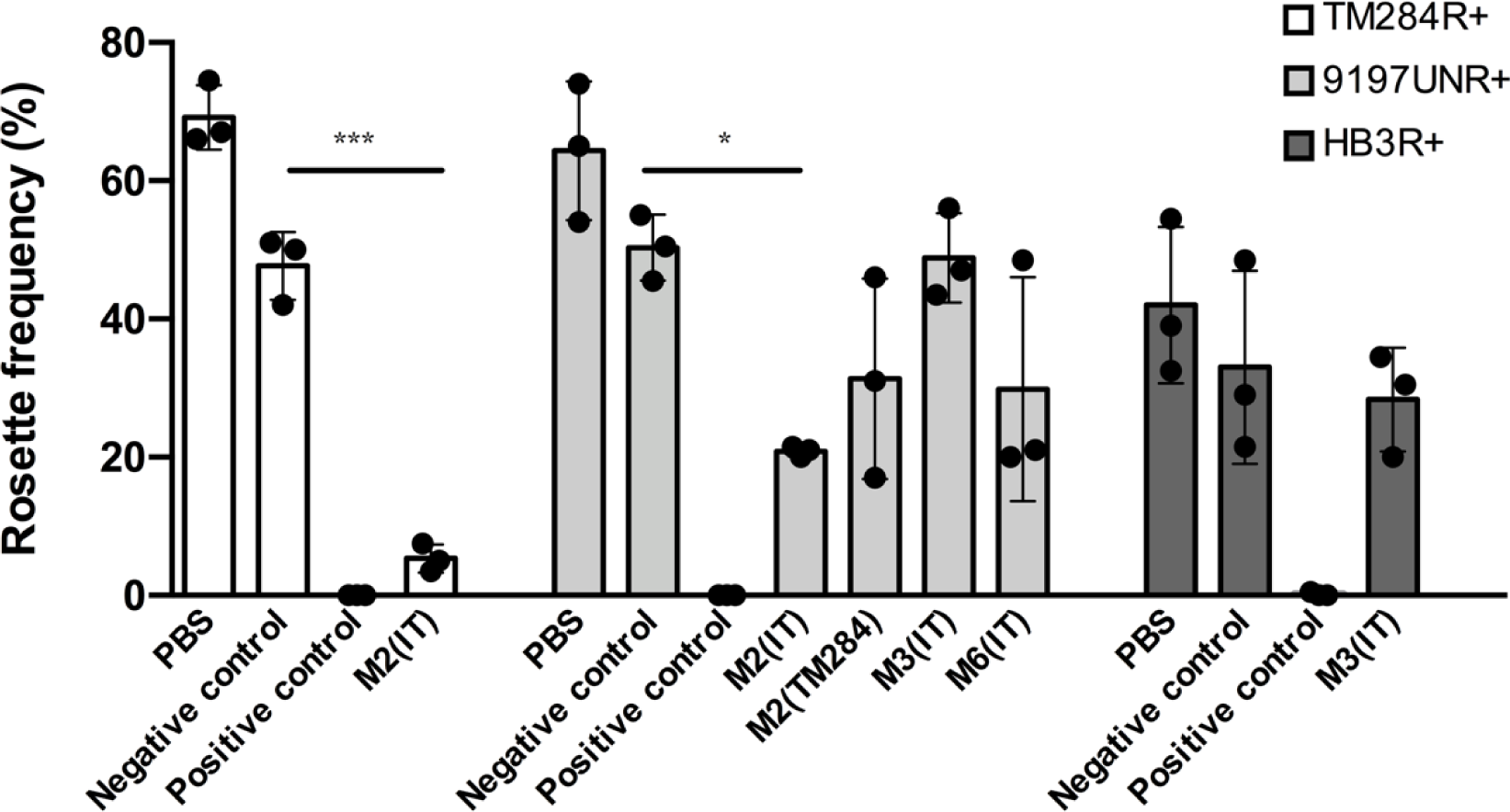
Rosette disrupting activity of eluates. Rosette frequency of parasite lines TM284R+, 9197UNR+ and HB3R+ (indicated in key) incubated in the presence of a negative control eluate that does not recognise mature infected erythrocytes, a positive control (antisera to the relevant DBLα PfEMP1 domain or heparin) showed surface recognition against the tested parasite line. The plasma and eluting line for each eluate is indicated on the X axis. Results from three independent experiments are shown as individual data points. Bar heights show the mean of the three experiments, and error bars indicate the SD. Significant differences between the negative control and tested eluate by student’s t-test or one way ANOVA are shown, ***P<0.001, *P<0.05. All differences between the negative and positive controls were statistically significant (P<0.0001).

## Discussion

Here we have shown that both adults and children living in *P. falciparum* endemic regions have naturally acquired IgG responses to allopatric rosetting parasite strains. In adults, these responses were common, with seroprevalence ranging from 58%-93%, whereas in children the seroprevalence was lower, ranging from 5%-30%. These findings build on previous studies that have described high seroprevalence in adults of antibodies that recognise rosetting parasite strains and the age-related build-up of anti-rosetting immunity [4,6,20,51,52]. The results imply that *P. falciparum* isolates that share antigenic determinants with the culture-adapted rosetting strains used in this study circulate in the regions from which the plasma samples were derived, and that exposure to such isolates induces antibodies in the majority of the adult population. Given the diverse geographical origins of the plasma and parasite strains tested, this is in keeping with the existence of globally conserved epitopes on the surface of rosetting infected erythrocytes [7].

Using the eluted antibody technique, we went on to demonstrate the presence of strain-transcending antibodies in both adults and children that recognise the infected erythrocyte surface of genetically distinct rosetting strains. The use of acid elution to show the existence of strain-transcending antibodies against epitopes on live infected erythrocytes was first reported by Marsh and Howard in the 1980s [19]. Surprisingly, in the four decades since then, this result has not to our knowledge been replicated until the work reported here. A more recent study by Tan et al. [24] used the mixed agglutination technique to show the existence of rare strain-transcending antibodies in Kenyan adults, which target some RIFIN family members. Mixed agglutination relies on the ability of bivalent IgG to cause mixed colour agglutinates if surface epitopes are shared between genetically distinct infected erythrocytes pre-stained with different colours [53]. Another previous study using mixed agglutination sought to identify strain-transcending antibodies to three rosetting *P. falciparum* strains, Palo Alto VarO, IT4 R29 and 3D7 PF13 in Senegalese sera, but found only variant-specific responses [6]. However, a study in Pakistan identified agglutinating and rosette-disrupting responses to heterologous parasites in convalescent serum from children recovering from malaria, suggesting the presence of strain-transcending responses to rosetting strains [54]. Our results unequivocally demonstrate that human IgG eluted from the infected erythrocyte surface of one rosetting strain can recognise multiple heterologous rosetting strains, and therefore prove the existence of naturally acquired strain-transcending antibodies to rosetting infected erythrocytes in humans. This in turn implies the presence of conserved epitopes on infected erythrocyte surface antigens displayed by rosetting parasite strains.

Current evidence for the commonness of strain-transcending antibodies against *P. falciparum* VSAs is contradictory. Most previous studies using mixed agglutination to look for strain-transcending antibodies to the surface of infected erythrocytes indicate that strain-transcending activity is rarely detected in plasma samples from malaria endemic regions [24,53,55]. Similarly, an extensive competition ELISA suggested that strain-transcending activity to PfEMP1 was very rare, only being detected in 1% of the 1245 heterologous protein combinations tested [56]. In contrast, one study detected frequent mixed strain agglutinates in the presence of plasma from malaria-exposed Indian donors [57], and serological studies have demonstrated recognition of allopatric isolates, with which the donors could never have been infected [16,19]. Furthermore, people who have only ever been exposed to a single infecting genotype, such as tourists returning from malaria endemic areas, or participants in experimental malaria challenge studies, have been reported to develop antibodies to heterologous *P. falciparum* strains, or recombinant PfEMP1 domains from heterologous infected erythrocytes [58–60].

Some of the contradictory findings from prior studies may be explained if epitopes are only shared between functionally related PfEMP1 variants. In such a scenario, the choice of parasite strains examined in any study is crucial, and strain-transcending activity will only be detected if parasite strains expressing functionally related subsets of PfEMP1 are used. Prior studies have shown strain-transcending activity of human mAbs against the VAR2CSA PfEMP1 variant, which is responsible for infected erythrocyte adhesion to Chondroitin Sulfate A (CSA) in the placenta during pregnancy malaria [21,22]. These mAbs recognise the infected erythrocyte surface of VAR2CSA-expressing culture-adapted parasite strains and clinical isolates, but do not recognise parasite isolates expressing other PfEMP1 variant types. Our own prior work shows that antibodies generated in rabbits against rosette-mediating PfEMP1 variants have strain-transcending activity against lab strains and clinical isolates, but only against parasites with same dual rosetting IgM Fc-binding phenotype [7]. Strain-transcending activity has also been described for antibodies against parasite strains showing adhesion to ICAM-1 [61] and EPCR [62], although much of the data for these functionally related subsets is based on recombinant protein assays. This is important because for PfEMP1, recognition and cross-reactivity of antibodies in ELISA-type assays using recombinant protein does not necessarily reflect recognition of native PfEMP1 on the infected erythrocyte surface [6][7][63]. Hence, flow cytometry, agglutination assays or other functional experiments with live infected erythrocytes are needed to prove the presence of strain-transcending antibodies against native surface antigens.

Although the target of the strain-transcending antibodies described here is unknown and will require further investigation, we provide preliminary evidence that the rosette-mediating adhesin PfEMP1 may be the target of antibodies in at least one of the eluates, because the eluted IgG was able to disrupt the rosettes in two heterologous parasite strains. It is possible that the targets of the strain-transcending antibodies identified in this work are antigens other than PfEMP1. Recently, Tan and colleagues reported the existence of broadly strain-transcending antibodies to the surface of infected erythrocytes which recognised RIFINs [24]. However, the strain-transcending activity observed here is unlikely to be due to the LAIR-1 insertions described by Tan et al., as such antibodies were rare, being found in only 3 out of 557 Kenyan adult plasma examined [24].

The work presented here has several limitations. In the eluted antibody experiments, strain-transcending activity could be missed due to incomplete elution of IgG from the surface of the adsorbing infected erythrocytes, which occurred commonly (data not shown). Furthermore, low concentrations of strain-transcending antibodies in plasma would be hard to detect using our methods, and our approach does not identify the target(s) of strain-transcending antibodies. This study does, however, confirm the biological relevance of strain-transcending antibodies, and by using live infected erythrocytes rather than recombinant proteins, it avoids the problems associated with serological assays in which recognition of recombinant PfEMP1 domains does not necessarily indicate recognition of native protein [63]. Despite the limitations of our study, the finding of high seroprevalence against allopatric rosetting isolates in adults from malaria-endemic regions, along with the demonstration of relatively common strain-transcending antibodies to the rosetting infected erythrocyte surface, are encouraging findings that warrant further investigation. Future experiments will aim to identify the targets of strain-transcending antibodies on the infected erythrocyte surface, and to characterise the conserved epitopes that are recognised. Identification of cryptic conserved infected erythrocyte surface epitopes may be a useful approach for the development of interventions against severe malaria [64].

## Acknowledgements

We are grateful to the donors from Ghana and the previous studies (Kenya, Malawi, Mali, and Papua New Guinea) who donated the plasma samples used in this work. We thank Dr. Eugene Martey (Komfo Anokye Teaching Hospital) and Amos Kotey (Agogo Presbyterian Hospital) for their help in the collection of the Ghana plasma samples. We also thank Professor Kevin Marsh and Professor Stephen Rogerson and colleagues for help in collection of samples in prior studies and for comments on the manuscript. This research was funded in part by the Wellcome Trust [grant number 204052/Z/16/Z] and by the Darwin Trust of Edinburgh (PhD studentship to Yvonne Azasi). For the purpose of open access, the author has applied a CC BY public copyright licence to any Author Accepted Manuscript version arising from this submission.

## Supplementary figures

**Figure S1.**
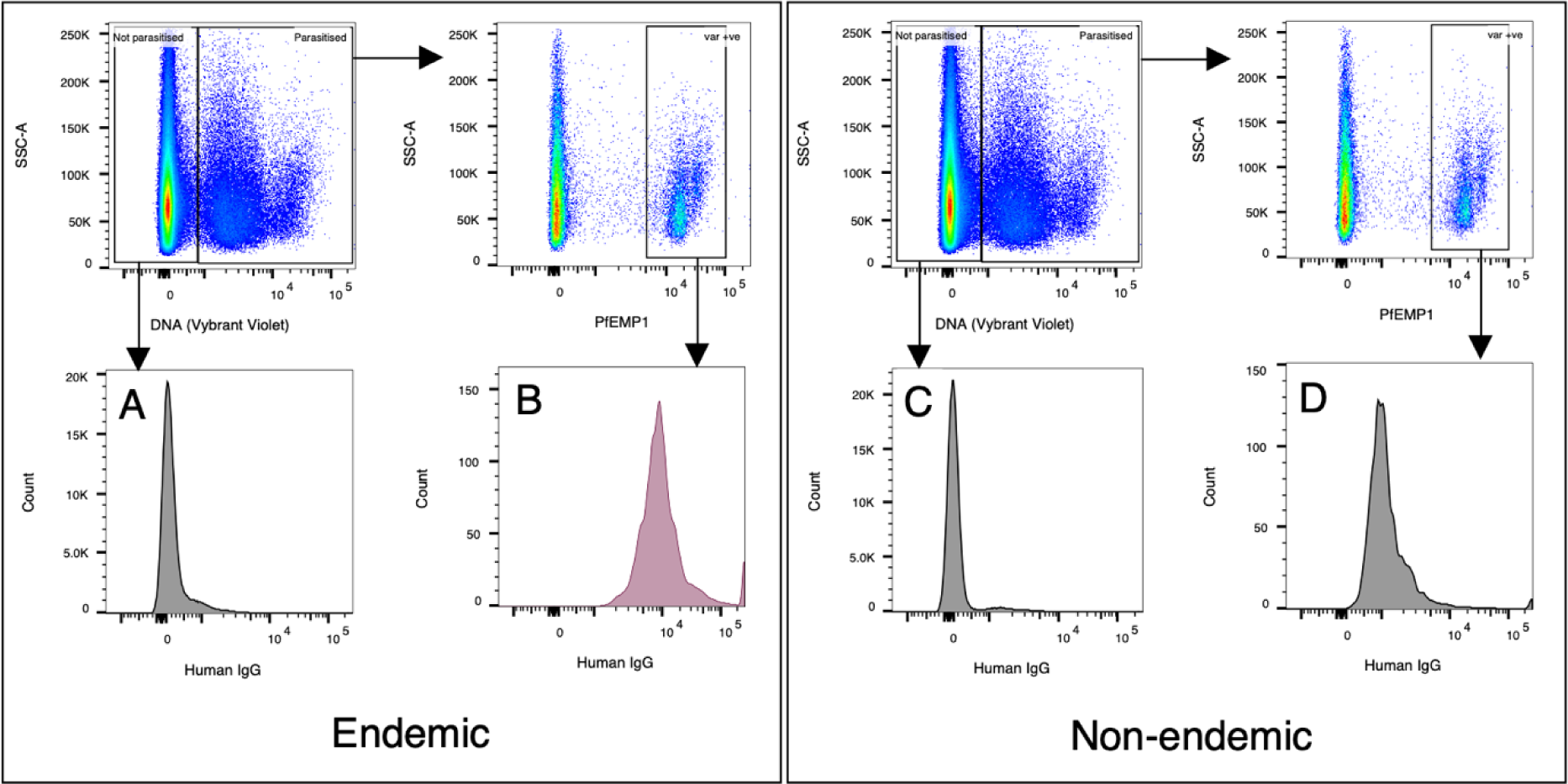
Correction of the Alexa Fluor^TM^ 488 median fluorescence intensity. Left panel, an example of gating and human IgG staining for a *P. falciparum* culture incubated with plasma (1/10 dilution) from a malaria-exposed individual (“Endemic”). Right panel, an example of gating and human IgG staining for the same *P. falciparum* culture incubated with 1/10 dilution of the negative control European serum pool (“Non-endemic”). The specific human IgG response is quantified by subtracting the Alexa Fluor^TM^ 488 MFI of the uninfected erythrocyte population (Vybrant^TM^ Dyecycle^TM^ Violet negative) (A) from the Alexa Fluor^TM^ 488 MFI of the infected erythrocyte population expressing the PfEMP1 variant of interest (B), which corrects for the presence of any anti-erythrocyte antibodies in the “endemic” sample. This is further corrected for any background staining observed with non-endemic negative control sera on infected erythrocytes expressing the PfEMP1 variant of interest (D), minus the MFI of uninfected erythrocytes with the negative control pool (C). Overall, this gives the cMFI = (B - A) - (D - C) [43]. The signal from the test plasma is shown in pink and the negative control population in grey. DNA staining was with 1/2500 dilution of Vybrant^TM^ Dyecycle^TM^ Violet (upper left panels). PfEMP1 was detected with 20μg/ml rabbit polyclonal IgG against the NTS-DBLα domain of the PfEMP1 variant of interest and an Alexa Fluor^TM^ 647-conjugated anti-rabbit IgG secondary antibody at 1/1000 (upper right panels). An Alexa Fluor^TM^ 488-conjugated anti-human IgG (gamma chain) antibody was used to detect human IgG at 1/1000 dilution (lower panels).

**Figure S2.**
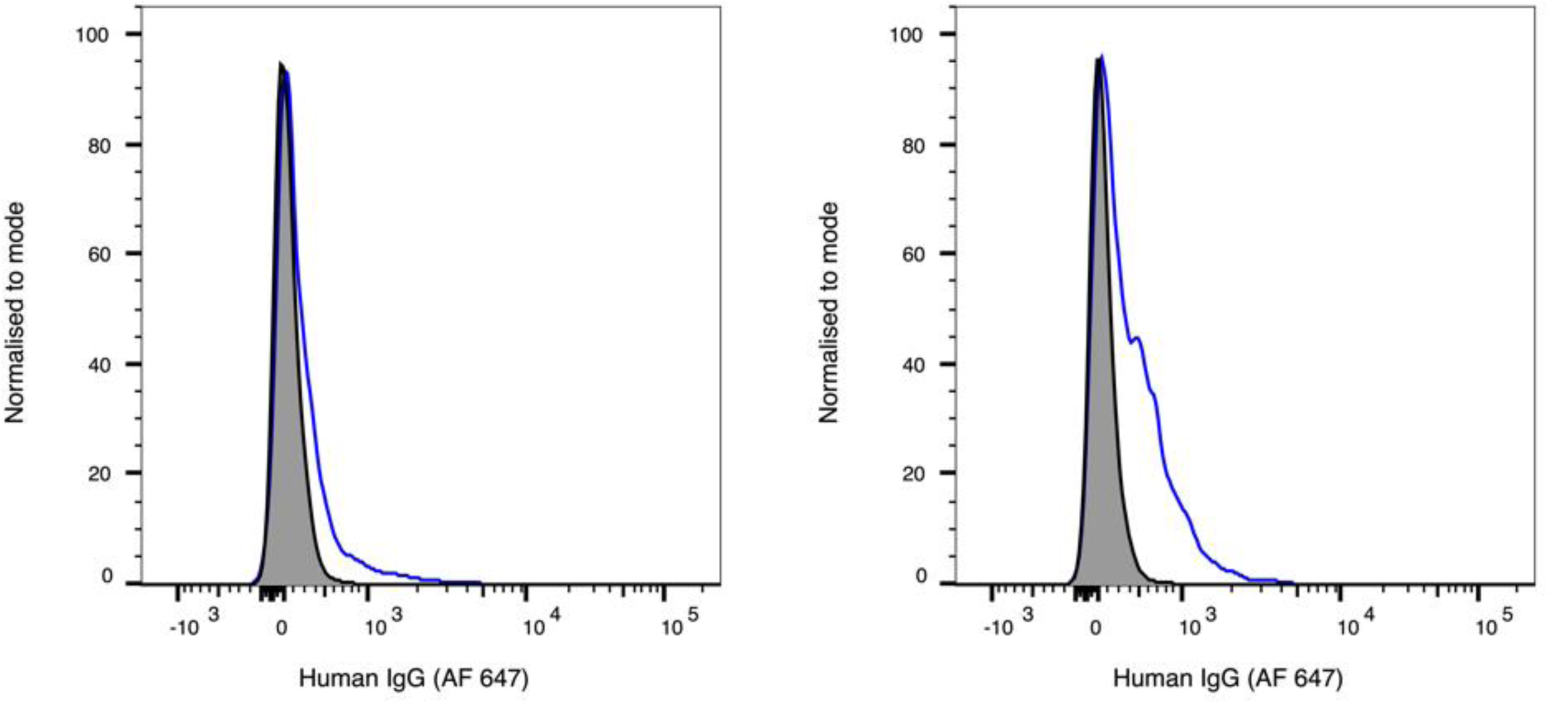
Sample above and below the minimum corrected fluorescence intensity positivity cut-off of 100 fu. Fluorescence intensity histograms of mature infected erythrocytes (blue, gated as described in Figure 5) with cMFI values of 50 (left), considered negative, and 155 (right) considered positive compared to a negative control (grey). Plasma was incubated at 1/10 dilution and human IgG was detected on the surface of infected erythrocytes with an Alexa Fluor^TM^ 647-conjugated anti-human IgG antibody at a 1/1000 dilution.

**Figure S3.**
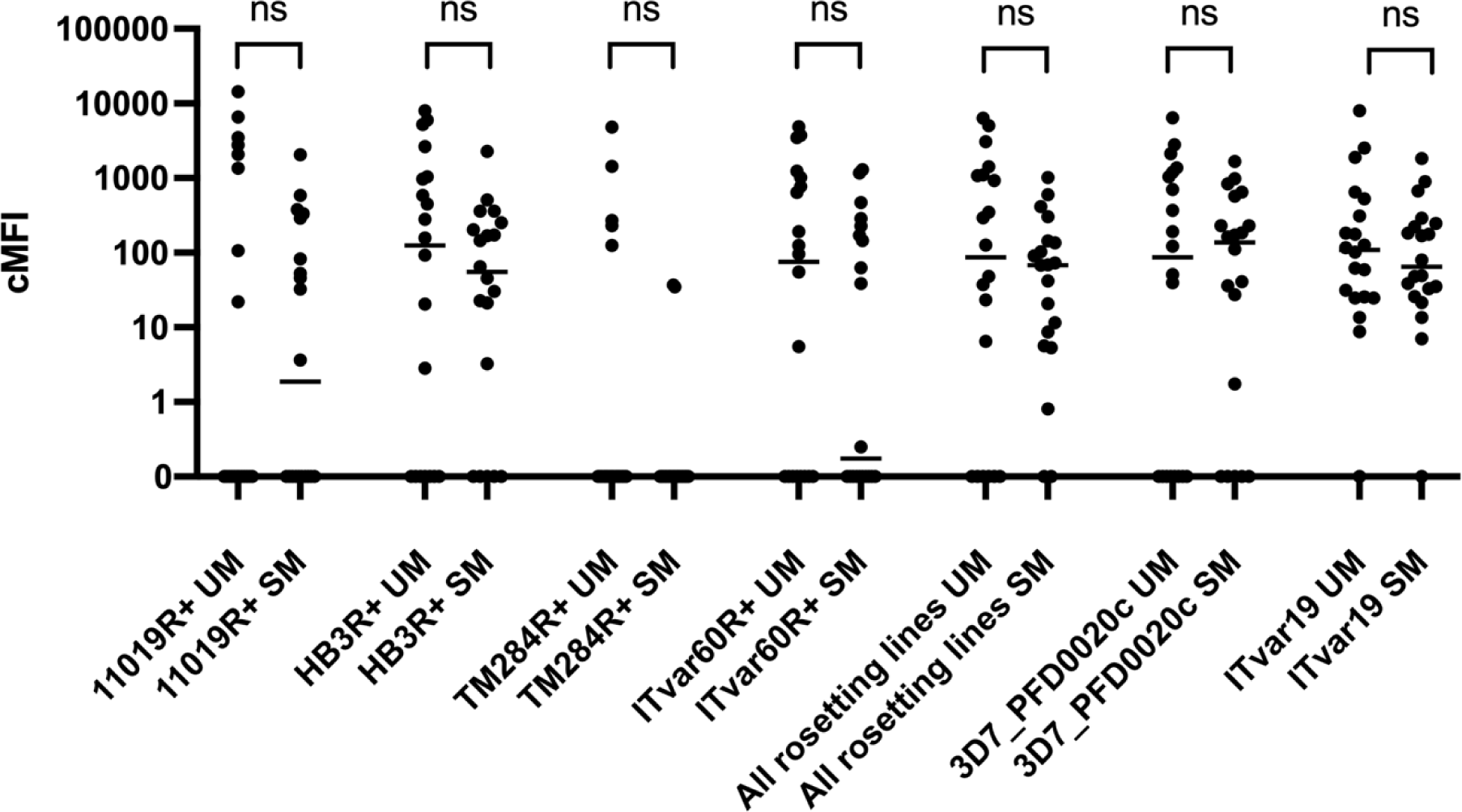
Strength of infected erythrocyte recognition by disease category. Corrected median fluorescence intensities (cMFI) of rosetting PfEMP1 variant*-*expressing infected erythrocytes stained by indirect immunofluorescence for recognition with childrens’ plasma, presented by parasite strain and disease category. Each dot represents one plasma sample, and gives the mean cMFI of two independent experiments for each sample. A horizontal line indicates the median for each category. Where the median is not visible, this is because it is 0 and so overlays the x-axis. cMFIs from individuals with uncomplicated and severe malaria were compared using the Mann-Whitney U test and the results indicated. UM, uncomplicated malaria, SM, severe malaria, ns, not significant. “All rosetting lines” shows the mean cMFI of all four rosetting parasite strains for each individual.

## References

[1] Smith JD, Rowe JA, Higgins MK, Lavstsen T. Malaria’s deadly grip: cytoadhesion of Plasmodium falciparum-infected erythrocytes. Cell Microbiol 2013;15:1976–83. 10.1111/cmi.12183.

[2] Wahlgren M, Goel S, Akhouri RR. Variant surface antigens of Plasmodium falciparum and their roles in severe malaria. Nat Rev Microbiol 2017;15:479–91. 10.1038/nrmicro.2017.47.

[3] Rowe JA, Moulds JM, Newbold CI, Miller LH. P. falciparum rosetting mediated by a parasite-variant erythrocyte membrane protein and complement-receptor 1. Nature 1997;388:292–5. 10.1038/40888.

[4] Vigan-Womas I, Guillotte M, Le Scanf C, Igonet S, Petres S, Juillerat A, et al. An in vivo and in vitro model of Plasmodium falciparum rosetting and autoagglutination mediated by varO, a group A var gene encoding a frequent serotype. Infect Immun 2008;76:5565–80. 10.1128/IAI.00901-08.

[5] Albrecht L, Moll K, Blomqvist K, Normark J, Chen Q, Wahlgren M. var gene transcription and PfEMP1 expression in the rosetting and cytoadhesive Plasmodium falciparum clone FCR3S1.2. Malar J 2011;10:17. 10.1186/1475-2875-10-17.

[6] Vigan-Womas I, Guillotte M, Juillerat A, Vallieres C, Lewit-Bentley A, Tall A, et al. Allelic diversity of the Plasmodium falciparum erythrocyte membrane protein 1 entails variant-specific red cell surface epitopes. PLoS ONE 2011;6:e16544. 10.1371/journal.pone.0016544.

[7] Ghumra A, Semblat J-P, Ataide R, Kifude C, Adams Y, Claessens A, et al. Induction of strain-transcending antibodies against Group A PfEMP1 surface antigens from virulent malaria parasites. PLoS Pathog 2012;8:e1002665. 10.1371/journal.ppat.1002665.

[8] Goel S, Palmkvist M, Moll K, Joannin N, Lara P, Akhouri RR, et al. RIFINs are adhesins implicated in severe Plasmodium falciparum malaria. Nat Med 2015;21:314–7. 10.1038/nm.3812.

[9] Niang M, Bei AK, Madnani KG, Pelly S, Dankwa S, Kanjee U, et al. STEVOR is a Plasmodium falciparum erythrocyte binding protein that mediates merozoite invasion and rosetting. Cell Host Microbe 2014;16:81–93. 10.1016/j.chom.2014.06.004.

[10] Kaul DK, Roth EF, Nagel RL, Howard RJ, Handunnetti SM. Rosetting of Plasmodium falciparum-infected red blood cells with uninfected red blood cells enhances microvascular obstruction under flow conditions. Blood 1991;78:812–9.

[11] Carlson J, Helmby H, Hill AV, Brewster D, Greenwood BM, Wahlgren M. Human cerebral malaria: association with erythrocyte rosetting and lack of anti-rosetting antibodies. Lancet 1990;336:1457–60. 10.1016/0140-6736(90)93174-n.

[12] Treutiger CJ, Hedlund I, Helmby H, Carlson J, Jepson A, Twumasi P, et al. Rosette formation in Plasmodium falciparum isolates and anti-rosette activity of sera from Gambians with cerebral or uncomplicated malaria. Am J Trop Med Hyg 1992;46:503–10. 10.4269/ajtmh.1992.46.503.

[13] Rowe A, Obeiro J, Newbold CI, Marsh K. Plasmodium falciparum rosetting is associated with malaria severity in Kenya. Infect Immun 1995;63:2323–6. 10.1128/iai.63.6.2323-2326.1995.

[14] Doumbo OK, Thera MA, Koné AK, Raza A, Tempest LJ, Lyke KE, et al. High levels of Plasmodium falciparum rosetting in all clinical forms of severe malaria in African children. Am J Trop Med Hyg 2009;81:987–93. 10.4269/ajtmh.2009.09-0406.

[15] Bull PC, Marsh K. The role of antibodies to Plasmodium falciparum-infected-erythrocyte surface antigens in naturally acquired immunity to malaria. Trends Microbiol 2002;10:55–8. 10.1016/s0966-842x(01)02278-8.

[16] Bull PC, Abdi AI. The role of PfEMP1 as targets of naturally acquired immunity to childhood malaria: prospects for a vaccine. Parasitology 2016;143:171–86. 10.1017/S0031182015001274.

[17] Rask TS, Hansen DA, Theander TG, Gorm Pedersen A, Lavstsen T. Plasmodium falciparum erythrocyte membrane protein 1 diversity in seven genomes--divide and conquer. PLoS Comput Biol 2010;6. 10.1371/journal.pcbi.1000933.

[18] Gonçalves BP, Huang C-Y, Morrison R, Holte S, Kabyemela E, Prevots DR, et al. Parasite burden and severity of malaria in Tanzanian children. N Engl J Med 2014;370:1799–808. 10.1056/NEJMoa1303944.

[19] Marsh K, Howard RJ. Antigens induced on erythrocytes by P. falciparum: expression of diverse and conserved determinants. Science 1986;231:150–3. 10.1126/science.2417315.

[20] Barragan A, Kremsner PG, Weiss W, Wahlgren M, Carlson J. Age-related buildup of humoral immunity against epitopes for rosette formation and agglutination in African areas of malaria endemicity. Infect Immun 1998;66:4783–7.

[21] Barfod L, Bernasconi NL, Dahlbäck M, Jarrossay D, Andersen PH, Salanti A, et al. Human pregnancy-associated malaria-specific B cells target polymorphic, conformational epitopes in VAR2CSA. Mol Microbiol 2007;63:335–47. 10.1111/j.1365-2958.2006.05503.x.

[22] Barfod L, Dobrilovic T, Magistrado P, Khunrae P, Viwami F, Bruun J, et al. Chondroitin sulfate A-adhering Plasmodium falciparum-infected erythrocytes express functionally important antibody epitopes shared by multiple variants. J Immunol 2010;185:7553–61. 10.4049/jimmunol.1002390.

[23] Doritchamou JYA, Renn JP, Jenkins B, Mahamar A, Dicko A, Fried M, et al. A single full-length VAR2CSA ectodomain variant purifies broadly neutralizing antibodies against placental malaria isolates. ELife 2022;11. 10.7554/eLife.76264.

[24] Tan J, Pieper K, Piccoli L, Abdi A, Perez MF, Geiger R, et al. A LAIR1 insertion generates broadly reactive antibodies against malaria variant antigens. Nature 2016;529:105–9. 10.1038/nature16450.

[25] Scholander C, Treutiger CJ, Hultenby K, Wahlgren M. Novel fibrillar structure confers adhesive property to malaria-infected erythrocytes. Nat Med 1996;2:204–8. 10.1038/nm0296-204.

[26] Rowe JA, Shafi J, Kai OK, Marsh K, Raza A. Nonimmune IgM, but not IgG binds to the surface of Plasmodium falciparum-infected erythrocytes and correlates with rosetting and severe malaria. Am J Trop Med Hyg 2002;66:692–9. 10.4269/ajtmh.2002.66.692.

[27] Ghumra A, Semblat J-P, McIntosh RS, Raza A, Rasmussen IB, Braathen R, et al. Identification of residues in the Cmu4 domain of polymeric IgM essential for interaction with Plasmodium falciparum erythrocyte membrane protein 1 (PfEMP1). J Immunol 2008;181:1988–2000. 10.4049/jimmunol.181.3.1988.

[28] Moulds JM, Zimmerman PA, Doumbo OK, Kassambara L, Sagara I, Diallo DA, et al. Molecular identification of Knops blood group polymorphisms found in long homologous region D of complement receptor 1. Blood 2001;97:2879–85. 10.1182/blood.v97.9.2879.

[29] Duffy MF, Caragounis A, Noviyanti R, Kyriacou HM, Choong EK, Boysen K, et al. Transcribed var genes associated with placental malaria in Malawian women. Infect Immun 2006;74:4875–83. 10.1128/IAI.01978-05.

[30] Cockburn IA, Mackinnon MJ, O’Donnell A, Allen SJ, Moulds JM, Baisor M, et al. A human complement receptor 1 polymorphism that reduces Plasmodium falciparum rosetting confers protection against severe malaria. Proc Natl Acad Sci USA 2004;101:272–7. 10.1073/pnas.0305306101.

[31] Deans A-M, Lyke KE, Thera MA, Plowe CV, Koné A, Doumbo OK, et al. Low multiplication rates of African Plasmodium falciparum isolates and lack of association of multiplication rate and red blood cell selectivity with malaria virulence. Am J Trop Med Hyg 2006;74:554–63.

[32] Claessens A, Adams Y, Ghumra A, Lindergard G, Buchan CC, Andisi C, et al. A subset of group A-like var genes encodes the malaria parasite ligands for binding to human brain endothelial cells. Proc Natl Acad Sci USA 2012;109:E1772–81. 10.1073/pnas.1120461109.

[33] Handunnetti SM, Gilladoga AD, van Schravendijk MR, Nakamura K, Aikawa M, Howard RJ. Purification and in vitro selection of rosette-positive (R+) and rosette-negative (R-) phenotypes of knob-positive Plasmodium falciparum parasites. Am J Trop Med Hyg 1992;46:371–81. 10.4269/ajtmh.1992.46.371.

[34] Otto TD, Assefa SA, Böhme U, Sanders MJ, Kwiatkowski D, Pf3k consortium, et al. Evolutionary analysis of the most polymorphic gene family in falciparum malaria. [version 1; peer review: 1 approved, 2 approved with reservations]. Wellcome Open Res 2019;4:193. 10.12688/wellcomeopenres.15590.1.

[35] Robson KJH, Walliker D, Creasey A, McBride J, Beale G, Wilson RJM. Cross-contamination of Plasmodium cultures. Parasitology Today 1992;8:38–9. 10.1016/0169-4758(92)90075-D.

[36] Mu J, Awadalla P, Duan J, McGee KM, Joy DA, McVean GAT, et al. Recombination hotspots and population structure in Plasmodium falciparum. PLoS Biol 2005;3:e335. 10.1371/journal.pbio.0030335.

[37] Helmby H, Cavelier L, Pettersson U, Wahlgren M. Rosetting Plasmodium falciparum-infected erythrocytes express unique strain-specific antigens on their surface. Infect Immun 1993;61:284–8.

[38] Roberts DJ, Craig AG, Berendt AR, Pinches R, Nash G, Marsh K, et al. Rapid switching to multiple antigenic and adhesive phenotypes in malaria. Nature 1992;357:689–92. 10.1038/357689a0.

[39] Claessens A, Rowe JA. Selection of Plasmodium falciparum parasites for cytoadhesion to human brain endothelial cells. J Vis Exp 2012:e3122. 10.3791/3122.

[40] Bhasin VK, Trager W. Gametocyte-forming and non-gametocyte-forming clones of Plasmodium falciparum. Am J Trop Med Hyg 1984;33:534–7. 10.4269/ajtmh.1984.33.534.

[41] Kyriacou HM, Steen KE, Raza A, Arman M, Warimwe G, Bull PC, et al. In vitro inhibition of Plasmodium falciparum rosette formation by Curdlan sulfate. Antimicrob Agents Chemother 2007;51:1321–6. 10.1128/AAC.01216-06.

[42] Geiman QM, Meagher MJ. Susceptibility of a New World monkey to Plasmodium falciparum from man. Nature 1967;215:437–9. 10.1038/215437a0.

[43] Williams TN, Newbold CI. Reevaluation of flow cytometry for investigating antibody binding to the surface of Plasmodium falciparum trophozoite-infected red blood cells. Cytometry A 2003;56:96–103. 10.1002/cyto.a.10088.

[44] Moll K, Ljungström I, Perlmann H, Scherf A, Wahlgren M, editors. Methods In Malaria Research. 6th ed. EVIMalaR; 2013.

[45] Ch’ng J-H, Moll K, Quintana MDP, Chan SCL, Masters E, Moles E, et al. Rosette-Disrupting Effect of an Anti-Plasmodial Compound for the Potential Treatment of Plasmodium falciparum Malaria Complications. Sci Rep 2016;6:29317. 10.1038/srep29317.

[46] Peters JM, Fowler EV, Krause DR, Cheng Q, Gatton ML. Differential changes in Plasmodium falciparum var transcription during adaptation to culture. J Infect Dis 2007;195:748–55. 10.1086/511436.

[47] Bull PC, Lowe BS, Kortok M, Marsh K. Antibody recognition of Plasmodium falciparum erythrocyte surface antigens in Kenya: evidence for rare and prevalent variants. Infect Immun 1999;67:733–9.

[48] Bull PC, Kortok M, Kai O, Ndungu F, Ross A, Lowe BS, et al. Plasmodium falciparum-infected erythrocytes: agglutination by diverse Kenyan plasma is associated with severe disease and young host age. J Infect Dis 2000;182:252–9. 10.1086/315652.

[49] Nielsen MA, Staalsoe T, Kurtzhals JAL, Goka BQ, Dodoo D, Alifrangis M, et al. Plasmodium falciparum variant surface antigen expression varies between isolates causing severe and nonsevere malaria and is modified by acquired immunity. J Immunol 2002;168:3444–50. 10.4049/jimmunol.168.7.3444.

[50] Chan J-A, Boyle MJ, Moore KA, Reiling L, Lin Z, Hasang W, et al. Antibody Targets on the Surface of Plasmodium falciparum-Infected Erythrocytes That Are Associated With Immunity to Severe Malaria in Young Children. J Infect Dis 2019;219:819–28. 10.1093/infdis/jiy580.

[51] Vigan-Womas I, Lokossou A, Guillotte M, Juillerat A, Bentley G, Garcia A, et al. The humoral response to Plasmodium falciparum VarO rosetting variant and its association with protection against malaria in Beninese children. Malar J 2010;9:267. 10.1186/1475-2875-9-267.

[52] Quintana MDP, Ch’ng J-H, Moll K, Zandian A, Nilsson P, Idris ZM, et al. Antibodies in children with malaria to PfEMP1, RIFIN and SURFIN expressed at the Plasmodium falciparum parasitized red blood cell surface. Sci Rep 2018;8:3262. 10.1038/s41598-018-21026-4.

[53] Newbold CI, Pinches R, Roberts DJ, Marsh K. Plasmodium falciparum: the human agglutinating antibody response to the infected red cell surface is predominantly variant specific. Exp Parasitol 1992;75:281–92. 10.1016/0014-4894(92)90213-t.

[54] Iqbal J, Perlmann P, Berzins K. Serological diversity of antigens expressed on the surface of erythrocytes infected with Plasmodium falciparum. Trans R Soc Trop Med Hyg 1993;87:583–8. 10.1016/0035-9203(93)90097-a.

[55] Giha HA, Staalsoe T, Dodoo D, Elhassan IM, Roper C, Satti GM, et al. Overlapping antigenic repertoires of variant antigens expressed on the surface of erythrocytes infected by Plasmodium falciparum. Parasitology 1999;119 ( Pt 1):7–17.

[56] Joergensen L, Turner L, Magistrado P, Dahlbäck MA, Vestergaard LS, Lusingu JP, et al. Limited cross-reactivity among domains of the Plasmodium falciparum clone 3D7 erythrocyte membrane protein 1 family. Infect Immun 2006;74:6778–84. 10.1128/IAI.01187-06.

[57] Chattopadhyay R, Sharma A, Srivastava VK, Pati SS, Sharma SK, Das BS, et al. Plasmodium falciparum infection elicits both variant-specific and cross-reactive antibodies against variant surface antigens. Infect Immun 2003;71:597–604. 10.1128/iai.71.2.597-604.2003.

[58] Krause DR, Gatton ML, Frankland S, Eisen DP, Good MF, Tilley L, et al. Characterization of the antibody response against Plasmodium falciparum erythrocyte membrane protein 1 in human volunteers. Infect Immun 2007;75:5967–73. 10.1128/iai.00327-07.

[59] Elliott SR, Payne PD, Duffy MF, Byrne TJ, Tham W-H, Rogerson SJ, et al. Antibody recognition of heterologous variant surface antigens after a single Plasmodium falciparum infection in previously naive adults. Am J Trop Med Hyg 2007;76:860–4.

[60] Turner L, Wang CW, Lavstsen T, Mwakalinga SB, Sauerwein RW, Hermsen CC, et al. Antibodies against PfEMP1, RIFIN, MSP3 and GLURP are acquired during controlled Plasmodium falciparum malaria infections in naïve volunteers. PLoS ONE 2011;6:e29025. 10.1371/journal.pone.0029025.

[61] Bengtsson A, Joergensen L, Rask TS, Olsen RW, Andersen MA, Turner L, et al. A novel domain cassette identifies Plasmodium falciparum PfEMP1 proteins binding ICAM-1 and is a target of cross-reactive, adhesion-inhibitory antibodies. J Immunol 2013;190:240–9. 10.4049/jimmunol.1202578.

[62] Lau CKY, Turner L, Jespersen JS, Lowe ED, Petersen B, Wang CW, et al. Structural conservation despite huge sequence diversity allows EPCR binding by the PfEMP1 family implicated in severe childhood malaria. Cell Host Microbe 2015;17:118–29. 10.1016/j.chom.2014.11.007.

[63] Lopez-Perez M. mSphere of Influence: Going Native, or the Risk of Overreliance on Recombinant Antigens. MSphere 2020;5. 10.1128/mSphere.00224-20.

[64] Good MF, Yanow SK. Hiding in plain sight: an epitope-based strategy for a sub-unit malaria vaccine. Trends Parasitol 2023 Nov;39(11):929–935. doi:10.1016/j.pt.2023.08.006.

